# Natural variation in a short region of the *Acidovorax citrulli* type III-secreted effector AopW1 is associated with differences in cytotoxicity

**DOI:** 10.1101/2021.05.24.445476

**Authors:** Irene Jiménez-Guerrero, Monica Sonawane, Noam Eckshtain-Levi, Gustavo Mateus Da Silva, Francisco Pérez-Montaño, Meirav Leibman-Markus, Lianet Noda-Garcia, Maya Bar, Saul Burdman

**Affiliations:** Department of Plant Pathology and Microbiology, The Robert H. Smith Faculty of Agriculture, Food and Environment, The Hebrew University of Jerusalem, Rehovot, Israel; Department of Microbiology, University of Seville, Seville, Spain; Department of Plant Pathology and Weed Research, Agricultural Research Organization, The Volcani Institute, Bet Dagan, Israel

**Keywords:** *Acidovorax citrulli*, bacterial fruit blotch, type III secretion, effectors, actin, endosomes, EHD1

## Abstract

Bacterial fruit blotch (BFB) is a serious disease of melon and watermelon caused by the Gram-negative bacterium *Acidovorax citrulli*. The strains of the pathogen can be divided into two major genetic groups, I and II. While group I strains are strongly associated with melon, group II strains are more aggressive on watermelon. Like many pathogenic bacteria, *A. citrulli* secretes a variety of protein effectors to the host cell via the type III secretion system. In the present study, we characterized AopW1, an *A. citrulli* type III-secreted effector that shares similarity with the actin cytoskeleton-disrupting effector HopW1 of *Pseudomonas syringae* and with effectors from other plant-pathogenic bacterial species. *aopW1* is present in group I and II strains, encoding products of 485 amino acids. Although highly conserved in most of the sequence, AopW1 has a highly variable region (HVR) within amino acid positions 147 to 192, including 14 amino acid differences between groups. Here we show that group I AopW1 is more toxic to yeast and plant cells than group II AopW1, having a stronger actin filament disruption activity, and increased ability to reduce plant callose deposition. We demonstrate the role of some of these 14 amino acid positions in determining the phenotypic differences between the two versions of the effector. Moreover, cellular analyses revealed that in addition to the interaction with actin filaments, AopW1 is localized to the endoplasmic reticulum, chloroplasts, and early and recycling plant endosomes, with differences observed between the two AopW1 versions. Finally, we show that overexpression of the endosome-associated protein EHD1 that increases cellular recycling, attenuates the toxic effects exerted by AopW1 and increases defence responses. This study provides insights into the HopW1 family of bacterial effectors and their interactions with the plant cell and provides first evidence on the involvement of EHD1 in response to biotic stress.

## INTRODUCTION

Bacterial fruit blotch (BFB) of cucurbits is a devastating disease caused by the Gramnegative bacterium *Acidovorax citrulli*. Due to the impact of this disease on the cucurbit industry, and more specifically to melon and watermelon production, *A. citrulli* is considered as one of the most important plant-pathogenic species within the *Acidovorax* genus (Burdman and Walcott, 2012; Burdman and Walcott, 2018; Zhao and Walcott, 2018).

*Acidovorax citrulli* strains are divided into two major groups, I and II, that are readily distinguishable by carbon substrate utilization, whole cell fatty acid analysis, DNA-fingerprinting and comparative genome analysis (Burdman *et al*., 2005; Eckshtain-Levi *et al*., 2016; Feng *et al*., 2009; Walcott *et al*., 2000; Walcott *et al*., 2004). The two groups also differ in host preference. Group I strains have been isolated mainly from melon, but also from other non-watermelon cucurbit crops, whereas group II strains are more strongly associated with watermelon (Burdman *et al*., 2005; Walcott *et al*., 2000; Walcott *et al*., 2004). Recent experiments involving natural infection under field conditions have strengthened the differences in host preferential association between the two groups of strains (Zhao *et al*., 2020).

Plants utilize a defence barrier against pathogen infection via recognition of so-called pathogen- or microbe-associated molecular patterns (PAMPs and MAMPs, respectively). The perception process occurs through specific plant pattern recognition receptors (PRRs), that results in PAMP-triggered immunity (PTI), an array of defence responses that is able to arrest infection of most potential pathogens (Jones and Dangl, 2006; Tang *et al*., 2017). Conversely, pathogenic microbes are able to deliver sets of protein effectors into the host cell to promote virulence through alteration of the host cell metabolism and suppression of host defence responses (Feng and Zhou, 2012; Macho and Zipfel, 2015). *Acidovorax citrulli*, as similar to several other Gram-negative plant-pathogenic bacteria, utilizes a type III secretion system (T3SS) to deliver such effectors (Bahar and Burdman, 2010). On the other hand, plants evolved to recognize some effectors, initiating a strong defence response that is named effector-triggered immunity (ETI) and is often associated with a localized hypersensitive response (HR) that arrest pathogen infection (Jones and Dangl, 2006; Mansfield, 2009; Mudgett, 2005).

Eckshtain-Levi *et al*. (2014) carried out a comparative analysis of eleven genes encoding type III-secreted effectors (T3Es) of the group II model strain, AAC00-1. This study revealed that group I and II strains differ in their T3E arsenal. First, three of the eleven T3E genes of AAC00-1 were conserved in all tested group II strains, but were absent or had disrupted open reading frames (ORFs) in all tested group I strains. Second, while the other T3E genes were detected and shown to encode likely functional products in both groups, most of them were highly conserved within each group, and clustered separately across the two groups (Eckshtain-Levi *et al*., 2014). More recently, we used a multifaceted approach combining thorough sequence analysis, transcriptomics and machine learning, that led to the identification of 58 T3E genes of the group I model strain, M6 (Jiménez-Guerrero *et al*., 2020). This study ranked *A. citrulli* among the richest pathogenic bacteria in terms of T3E arsenal size, and confirmed the dissimilarity among group I and II strains of *A. citrulli* in terms of T3E cargo (Jiménez-Guerrero *et al*., 2020).

To date, only few studies were reported on the role of Aop (for *Acidovorax* outer proteins) T3Es in *A. citrulli*-plant interactions. AopP was shown to suppress reactive oxygen species burst and salicylic acid content and to significantly contribute to the virulence of a group II *A. citrulli* strain in watermelon. The authors suggested that AopP suppresses plant immunity by targeting ClWRKY6 in the plant nucleus (Zhang *et al*., 2020a). In addition, Zhang *et al*. (2020b) showed that the effector AopN locates to the cell membrane of *Nicotiana benthamiana* and induces a programmed cell death response in this plant.

We are particularly interested in effector AopW1 (*Acidovorax* outer protein W1). The genes encoding this effector are *APS58_3289* in the genome of the group I model strain M6, and *Aave_1548* in the genome of the group II model strain AAC00-1 (NCBI accessions CP02973.1 and NC_008752.1, respectively). The effector was named AopW1 due to its similarity with the *Pseudomonas syringae* pv. *maculicola* T3E, HopW1. Lee *et al*. (2008) showed that expression of *hopW1* in *P. syringae* pv. *tomato* triggers strong immunity in *N. benthamiana* and in *Arabidopsis thaliana* Ws, but promotes virulence in *A. thaliana* Col-0. The authors also showed that the C-terminal region of HopW1 is needed for triggering defence responses. Later, Kang *et al*. (2014) showed that filamentous actin (F-actin) is a major virulence target of HopW1. They showed that HopW1 interacts with and reduces actin filaments *in vitro* and *in planta*, disrupts the actin cytoskeleton, and affects actin-dependent cell biological processes that are critical for plant immunity like endocytosis and protein trafficking into vacuoles. Also, the C-terminal region of HopW1 was needed for these phenotypes (Kang *et al*., 2014).

As similar as several other *A. citrulli* T3Es, AopW1 is conserved within strains that belong to the same group but differ between the two groups. Specifically, group I and II versions of this effector are 485 amino acid (a.a.) long, and are highly conserved except for a hypervariable region (HVR) between a.a. positions 147 and 192, that shows 14 differences between the two versions (~30% dissimilarity) (Eckshtain-Levi *et al*., 2014). Traore *et al*. (2019) demonstrated that both group I and II AopW1 significantly contribute to the virulence of *A. citrulli* in melon and watermelon, respectively. Interestingly, in *Nicotiana tabacum*, group I AopW1 triggered a stronger HR than that exerted by group II AopW1, suggesting that the differences in the HVR between the two versions correlate with molecular function (Traore *et al*., 2019). Recently, we showed that in *A. citrulli* M6, expression of the *aopW1* gene is regulated by the T3SS transcriptional activator HrpX, and validated that AopW1 is translocated to the plant cell in a T3SS-dependent manner (Jiménez-Guerrero *et al*., 2020).

In the present study, we investigated the sequence-function relationship of AopW1. We asked whether differences in the HVR between group I and II AopW1 are determinants of their functional performance, including cytotoxicity, actin filament disruption ability, interaction with plant cell compartments, and ability to reduce callose deposition. Overall, our findings demonstrate that the group I version of AopW1 has increased toxicity as compared to group II AopW1. We also identified amino-acid residues in the HVR region that are likely critical for the activity of effectors belonging to the HopW1 family. Further, we found that the endosome-associated protein EHD1 attenuates AopW1-induced cytotoxicity. A general model of AopW1 function in the plant cell is proposed.

## RESULTS

### Heterologous expression in yeast reveals differences in cytotoxicity between group I and II versions of AopW1

To assess possible differences in cytotoxicity between group I and II AopW1, we used a yeast-based heterologous system that has been used to detect phenotypes of bacterial effectors (Salomon and Sessa, 2010; Siggers and Lesser, 2008). The *aopW1* open reading frames (ORFs) of strains M6 and 7a1 (belonging to groups I and II, respectively) were cloned into the yeast expression vector pGML10 and transferred to *Saccharomyces cerevisiae* BY4741 for heterologous expression. It is worth mentioning that the 7a1 *aopW1* ORF is 100% identical to that of the group II model strain AAC00-1 (Eckshtain-Levi *et al*., 2014).

Growth inhibition assays were carried out in inducing media (supplemented with 2% galactose and 1% raffinose), under regular conditions or with the addition of stressing compounds or conditions. It has been shown that combination of different stressors may increase yeast sensitivity thus aiding to detect effector-induced phenotypes (Salomon *et al*., 2011; Salomon *et al*., 2012; Siggers and Lesse, 2008). As stressors we used 7 mM caffeine, 0.5 M sodium chloride or 1 M sorbitol (Kuranda *et al*., 2006; Salomon *et al*., 2011; Yoon *et al*., 2003). Controls were yeast plated on repressing media (supplemented with 2% glucose).

Under regular conditions, group I AopW1 (AopW1-M6) exerted strong growth inhibition of yeast cells, while the group II version (AopW1-7a1) had a very slight effect (Figure 1A). The growth inhibition exerted by group I AopW1 was comparable to that observed for *P. syringae* HopW1 and slightly higher than that exerted by the HopW1-homologous effector from *Xanthomonas translucens* (Figure 1A; Supplementary Table S1). Growth inhibition by group I AopW1 was increased when combined with NaCl, while the growth inhibition effect of group II AopW1 was increased in combination with caffeine, NaCl or sorbitol, though at substantially lower levels than the effects exerted by the former (Supplementary Table S1).

**Figure 1.**
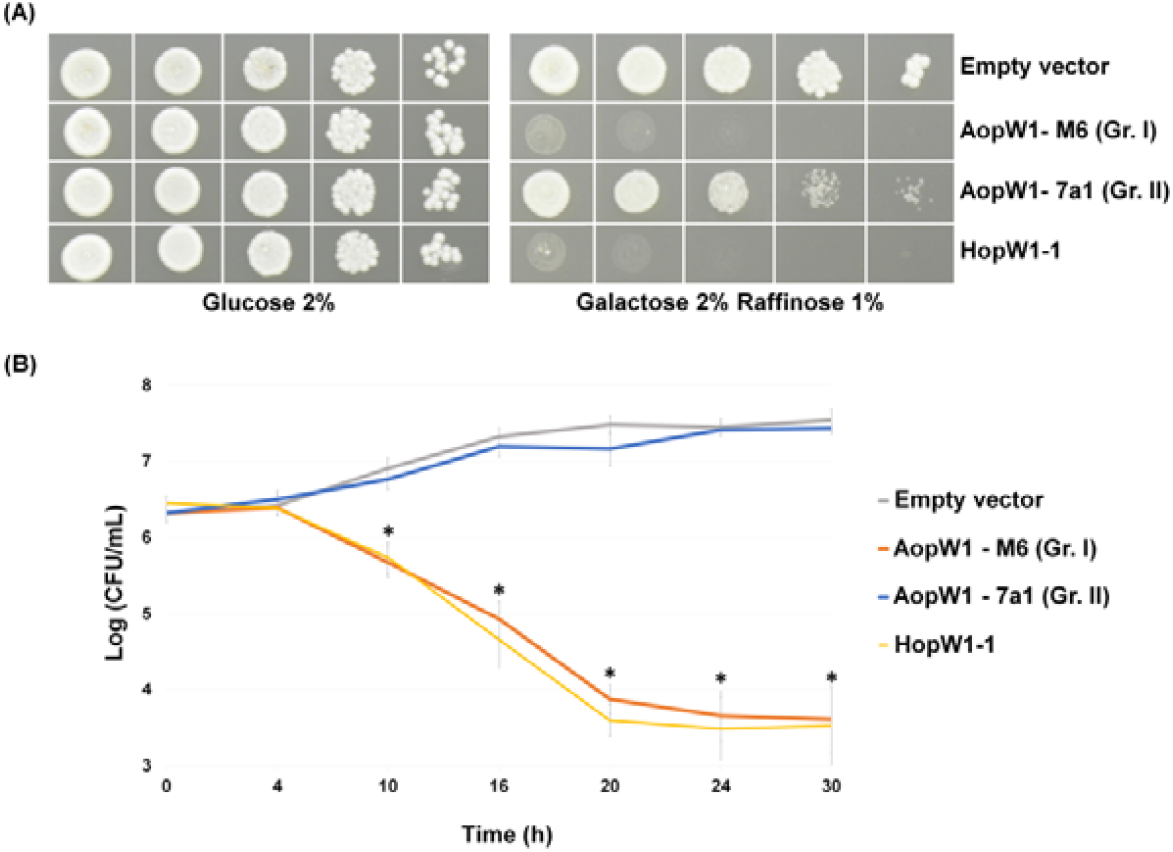
*Acidovorax citrulli* group I AopW1 is more toxic than group II AopW1 to yeast cells. **(A)** Yeast growth inhibition assays. The open reading frames (ORFs) of *aopW1* from *A. citrulli* strains M6 (group I) and 7a1 (group II), and *hopW1* from *Pseudomonas syringae* pv. *maculicola* ES4326 were cloned into pGML10 under the control of a galactose inducible promoter. The plasmids were transformed into *S. cerevisiae* BY4741. Serial dilutions of the yeast cultures were spotted in glucose (repressing) or in galactose/raffinose (inducing) media. Controls were yeast cells carrying pGML10 (empty vector). Pictures were taken after 3 days of growth at 28 °C and are representative of three independent experiments. **(B)** Cell viability assays of yeast expressing the aforementioned effectors. Yeast cells were grown in inducing medium, collected at different time points, 10-fold serially diluted, and spotted onto repressing medium to assess the number of viable cells. Data represent means and standard errors (SE) from one experiment out of three with similar results (with four replicates per treatment per time point). Asterisks indicate significant (p=0.05) differences compared to the empty vector each time point by Student t-test.

To assess whether yeast growth inhibition was due to cell death or growth arrest, we carried out yeast cell viability time-curves. A significant decrease of viable cells was detected in yeast expressing group I AopW1, in a similar manner as observed for expression of *P. syringae* HopW1. In contrast, the viability of cells expressing group II AopW1 did not significantly differ from controls carrying empty pGML10, demonstrating that this version of AopW1 does not exert a strong toxic effect on yeast (Figure 1B).

### Group I AopW1 strongly affects yeast F-actin organization *in vivo* and disrupts non-muscle F-actin *in vitro*

Actin is a highly abundant and conserved protein that is essential for the survival of most eukaryotic cells. It occurs in a globular monomeric (G-actin) stage or in a filamentous polymeric (F-actin) form (Pollard, 2016; Pollard *et al*., 2000). The actin cytoskeleton is a complex network of dynamic polymers, which has an important role in a wide range of cellular processes (Mishra *et al*., 2014; Pollard and Borisy, 2003). As aforementioned, Kang *et al*. (2014) showed that *P. syringae* HopW1 disrupts the actin cytoskeleton.

We studied the effects of AopW1 expression in yeast cells. Budding yeast cultures were stained with the actin stain TRITC-phalloidin 8 h after expression of group I or II AopW1. During budding, the actin filament network of yeast cells presents three specific structures: cortical actin patches, actin cables, and the actomyosin ring (Mishra *et al*., 2014; Supplementary Figure 1A). Yeast cells carrying empty pGML10 displayed normal formation of actin filaments and migration of actin patches to the daughter cells, and a similar picture was observed in cells expressing group II AopW1. In contrast, cells expressing group I AopW1 exhibited a strong disorganization of the actin structures, with actin patches seeming disorganized thorough the cell (Supplementary Figure 1B). Remarkably, in yeast cells expressing group I AopW1 we were able to detect substantially less formation of daughter cells than in the two other treatments.

To assess whether the effect caused by group I AopW1 is due to actin depolymerization activity, we carried out *in vitro* sedimentation assays with preassembled non-muscle F-actin. In these assays we used purified recombinant AopW1 from group I and II strains without the first 100 a.a. of their N-terminus (AopW1-M6_101-485_ and AopW1-7a1_101-485_, respectively) fused to a poly-histidine (His) tag in both extremes. The N-terminal part of AopW1 was not included because we were not able to express sufficient amounts of soluble, full-length AopW1. As shown below, the N-terminal 100 a.a. of this effector is not needed for cytotoxic activity in yeast. Expression and purification of the recombinant proteins were verified by SDS-PAGE and Western blot analysis (Supplementary Figure S2). Incubation of pre-assembled non-muscle Factin with poly-His-AopW1-M6_101-485_, but not with α-actinin (which binds to F-actin) or bovine serum albumin (BSA; which does not bind to F-actin), led to a reduction of Factin (pellet fraction; P), with a concomitant increase of G-actin (supernatant fraction; S) (Figure 2). AopW1-M6_101-485_ could be detected in the supernatant fractions by Western blot analysis using a poly-His tag monoclonal antibody (not shown). Recombinant group II AopW1 (poly-His-AopW1-7a1_101-485_) was able to induce actin depolymerization, but at substantially lower levels than the group I version (Figure 2).

**Figure 2.**
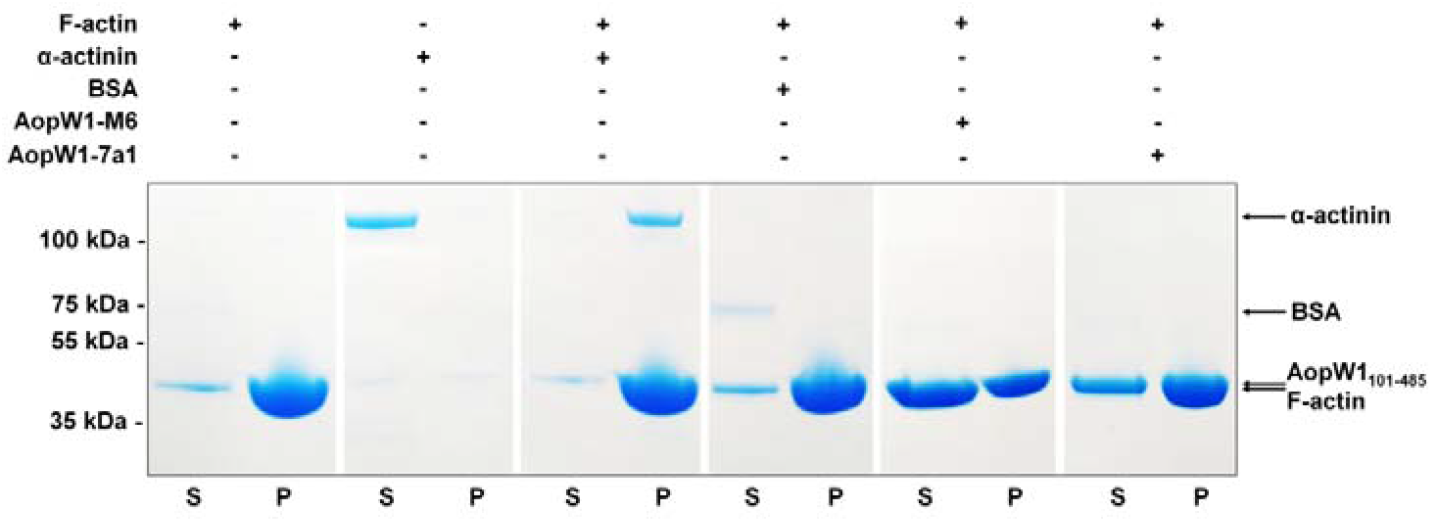
AopW1 disrupts F-actin *in vitro*. Preassembled F-actin (filamentous polymer) was incubated with 35 μg of AopW1101-485 from strains M6 (group I) or 7a1 (group II), 2 μM α-actinin or 2 μM BSA for 1.5 h at 24 °C and centrifuged at 150,000 *g* for 1.5 h. Samples were partitioned into supernatant (S) and pellet (P) fractions to separate G-actin (globular subunit) and F-actin, respectively. Proteins were separated by SDS-PAGE and stained with Coomassie blue. α-actinin was used as positive control for actin binding activity (presence in S or P when α-actinin was incubated alone or with F-actin, respectively), and as negative control for actin disruption activity (F-actin remained polymerized in P). BSA was used as negative control for actin binding and acting disrupting activities (presence in S when BSA was incubated with Factin; F-actin remained polymerized in P). Marker molecular masses (kDa) are shown in the left. The experiment was performed twice with similar results.

### Combined substitutions of specific amino acid residues alter yeast cytotoxicity of AopW1

T3Es sharing similarity with AopW1 occur in other plant-pathogenic bacteria. In addition to the aforementioned HopW1 from *P. syringae* and homologous effectors from *X. transluscens*, AopW1 homologs occur in other *Pseudomonas* spp., in *Erwinia mallotivora* and in other plant-pathogenic *Acidovorax* species (Figure 3A). As said, the *aopW1* gene is highly conserved among group I and II strains of *A. citrulli*, except for a 138-bp hypervariable region (HVR) located at nucleotide positions 439 to 576 (a.a. positions 147 to 192 in the encoded products). Of the 46 a.a. included in the HVR, fourteen are different between the group I and II versions of the effector; Figure 3B). Although group I and II versions of AopW1 share significantly higher levels of identity as compared with homologs from other bacteria, including in the HVR (Figure 3), of the aforementioned fourteen positions that distinguish between them, six are conserved between group I AopW1 and the homologous T3Es from *Pseudomonas, Xanthomonas* and *Erwinia* species. These are a.a. positions 154, 162, 167, 174, 177 and 189 (named positions 1 to 6, respectively, as indicated in the bottom of Figure 3B). Intriguingly, variation in the HVR also occurs among strains of *A. avenae*, with some variants resembling group I AopW1 and others resembling group II AopW1 (Figure 3B).

**Figure 3.**
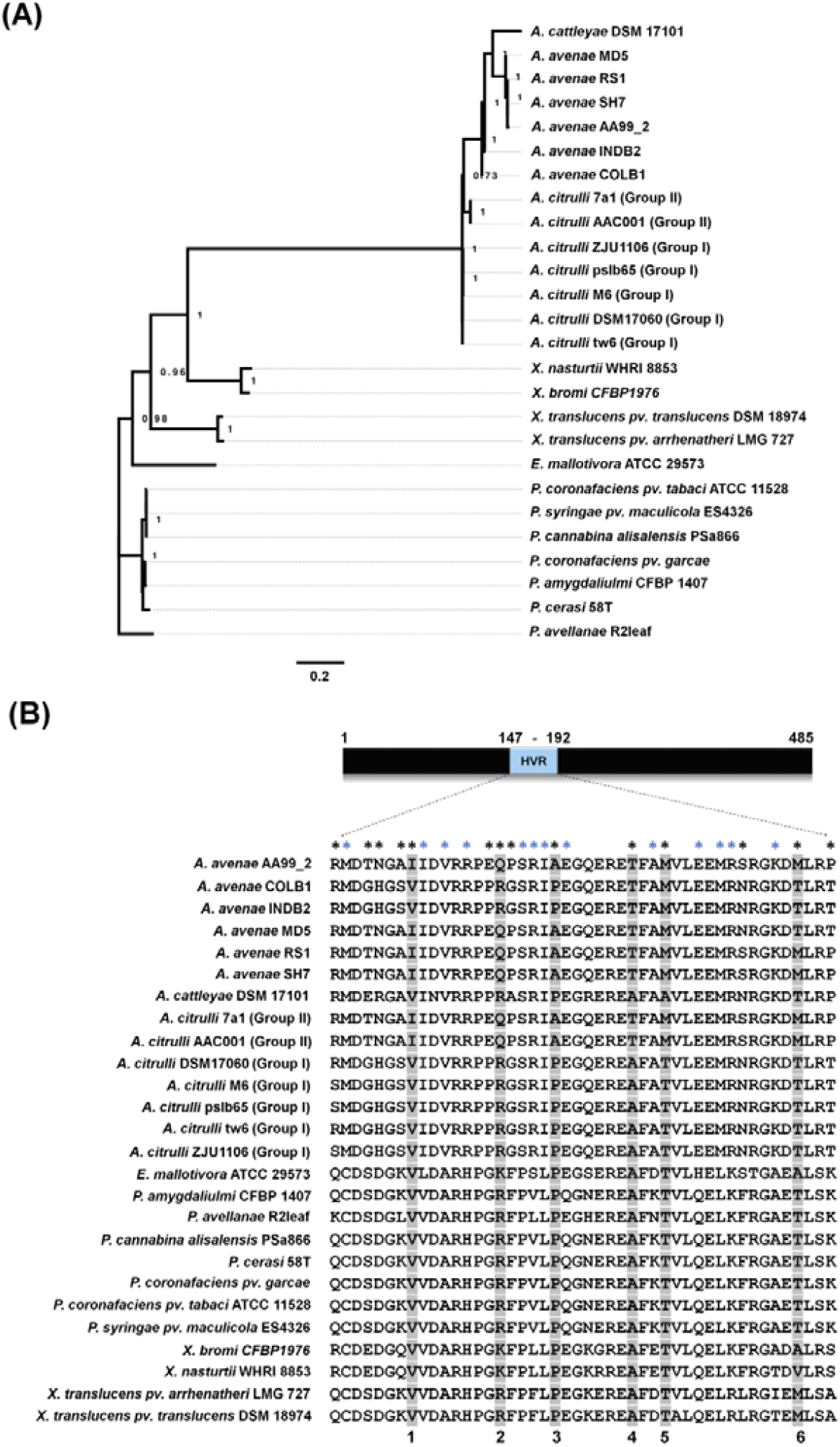
Comparison of AopW1 and homologous effectors. **(A)** AopW1 protein-based phylogeny from *Acidovorax, Pseudomonas, Xanthomonas* and *Erwinia* species. Sequences were aligned using MUSCLE and the phylogeny was reconstructed using maximum likelihood methods included in the SeaView program (ref: PMID 19854763). Numbers at nodes are the approximate likelihood ratio test supporting each branch. The scale bar shows amino acid (a.a.) substitutions per site. **(B)** Alignment of the hypervariable region (HVR) of *A. citrulli* AopW1 from group I and group II strains with corresponding regions from homologous effectors. The alignment corresponds to a.a. positions 147 to 192 of AopW1. The alignment was done using ClustalW software. Black asterisks indicate the 14 a.a. differences between the two versions of AopW1 in this region. Blue asterisks indicate the strongly conserved positions within *Acidovorax* sp. and that are different in other species. Grey shading and numbers at the bottom of the alignment indicate the six conserved a.a. positions between group I AopW1 and homologous effectors from other species.

We hypothesized that differences in the HVR, and particularly in the aforementioned six positions, are important determinants of the distinguished patterns of cytotoxicity between group I and II versions of AopW1. In this regard, it is worth mentioning that the AopW1 HVR aligns with part of the C-terminal region of *P. syringae* HopW1, which is required for actin cytoskeleton disruption (Kang *et al*., 2014).

To assess the importance of these six positions for AopW1 activity, we generated several constructs in which the group I and II *aopW1* ORFs were altered by site-directed mutagenesis to substitute each of these residues by the a.a. present in the other version (Supplementary Tables S2 and S3). We also generated group I and II AopW1 carrying multiple substitutions (Supplementary Table S3). All constructs were tested in yeast growth inhibition assays.

Most individual substitutions in the HVR did not lead to significant alterations in yeast growth inhibition ability in comparison with the corresponding wild-type versions (Supplementary Figure S3). The only exception was the AopW1 group II variant carrying an individual substitution in position 6 (M-189-T) that reduced cytotoxicity to the empty vector level (Supplementary Figure S3B). Besides the HVR, group I and II AopW1 differ in position 319 (leucine in group I; serine in group II). Swap substitutions in this position (L319S and S319L in group I and II AopW1, respectively; Supplementary Table S2) did not alter yeast growth inhibition ability as compared with the corresponding wild-type versions (not shown).

Characterization of combined substitutions in the background of group I AopW1 showed that only one variant, carrying three substitutions [1 (V-154-I) +2 (R-162-Q) +3 (P-167-A)], considerably reduced the cytotoxicity of the effector (Figure 4A). Surprisingly, adding substitutions in positions 4 (A-174-T) and/or 6 (T-189-M) to the above variant, abolished this effect, namely, led to a growth inhibition effect that was similar to that of group I AopW1 (Figure 4A).

**Figure 4.**
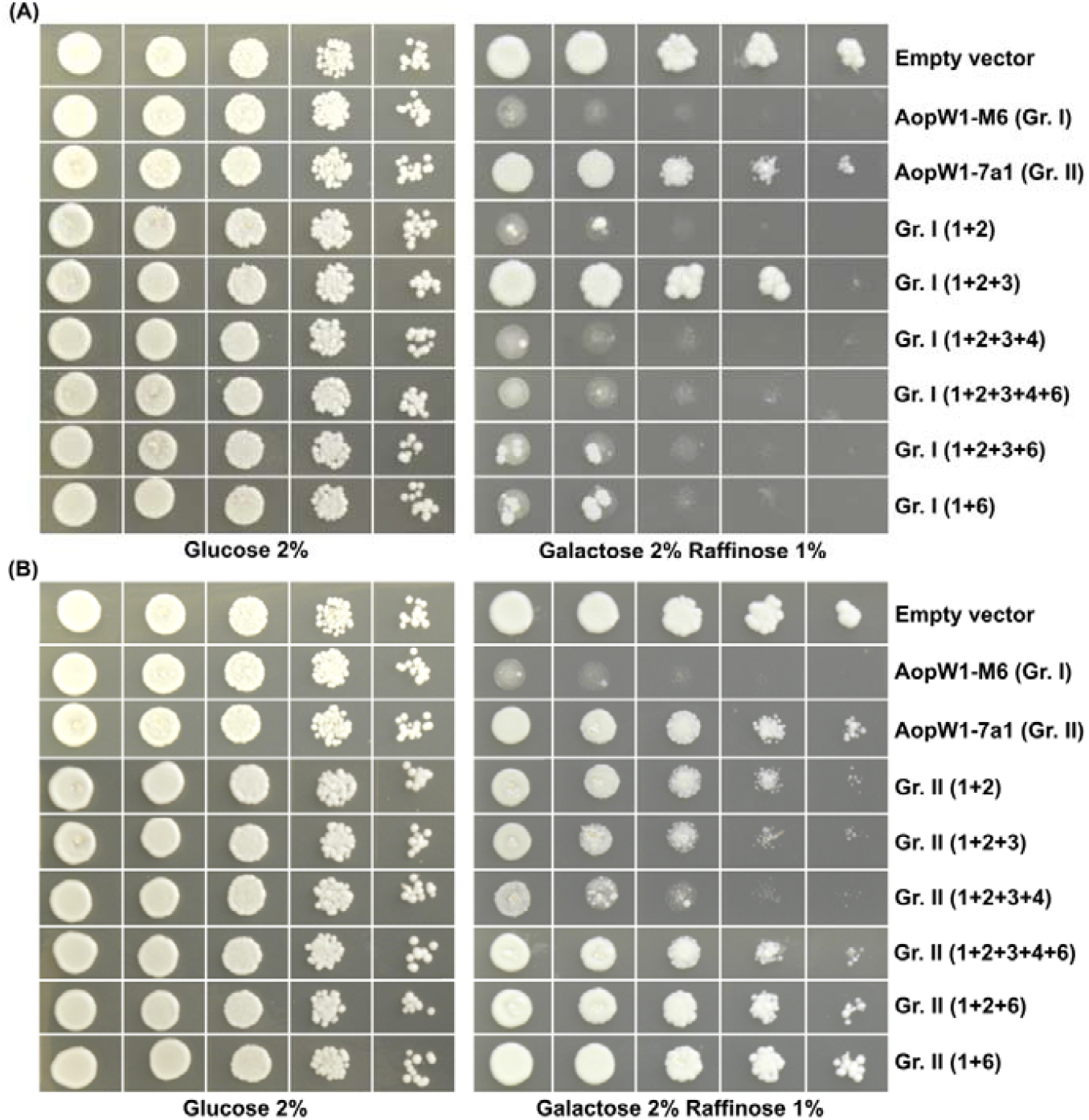
Effects of combined amino acid substitutions in the AopW1 HVR on yeast growth inhibition. Yeast growth inhibition assays of different variants of group I AopW1 **(A)** and group II AopW1 **(B)** carrying several combinations of a.a. substitutions in the six positions that are conserved among group I AopW1 and homologous effectors from other bacterial species (see Figure 4 and Supplementary Table S2 for variant details). *S. cerevisiae* BY4741 carrying the ORF of the *aopW1* variants in pGML10 were grown in glucose (repressing) and in galactose/raffinose (inducing) medium. Pictures were taken after 3 days of growth at 28 °C and are representative of three independent experiments.

In the case of group II AopW1, a variant carrying substitutions in positions 1 to 4 [1 (I-154-V) + 2 (Q-162-R) + 3 (A-167-P) + 4 (T-174-A)], had a substantially higher yeast growth inhibition ability as compared with the group II wild-type version (Figure 4B). However, adding a fifth substitution to this variant [6 (M-189-T)] abolished this effect. In addition, a combination of two substitutions [1 (I-154-V) + 2 (Q-162-R)] and a combination of three substitutions [1 (I-154-V) + 2 (Q-162-R) + 3 (A-167-P)] led to a subtle increment in the growth inhibition effect as compared with the group II wild-type version of AopW1 (Figure 4B). Overall, results from these experiments demonstrated that the AopW1 HVR is important for induction of cytotoxicity in yeast. These experiments also demonstrated the importance of the six positions that are conserved between group I AopW1 and most homologous effectors from other plant-pathogenic bacteria. However, it appears that the interactions among these positions are complex and not merely additive.

### The HVR and the C-terminal part of AopW1 are required for cytotoxic ability

To learn about the importance of the different parts of AopW1 for its activity, we generated pGML10 constructs carrying several N- and C-terminal truncated variants in the background of group I AopW1 (Supplementary Table S2; Figure 5). Growth inhibition assays with yeast cells carrying these constructs revealed that the N-terminal part of the effector (first 135 residues; AopW1-M6_Δ1-135_) is not important for yeast inhibition ability. However, removal of the next 10 residues (AopW1-M6_Δ1-145_) significantly reduced this ability. Further removal of 30 residues (AopW1-M6_Δ1-175_), which also removed part of the HVR, almost abolished yeast growth inhibition ability of AopW1. The C-terminal part was found to be critical for the cytotoxic ability of AopW1. While removal of the last 10 residues (AopW1-M6_Δ476-485_) did not affect AopW1 activity, removal of the last 25 residues (AopW1-M6_Δ461-485_) completely abolished it (Figure 5).

**Figure 5.**
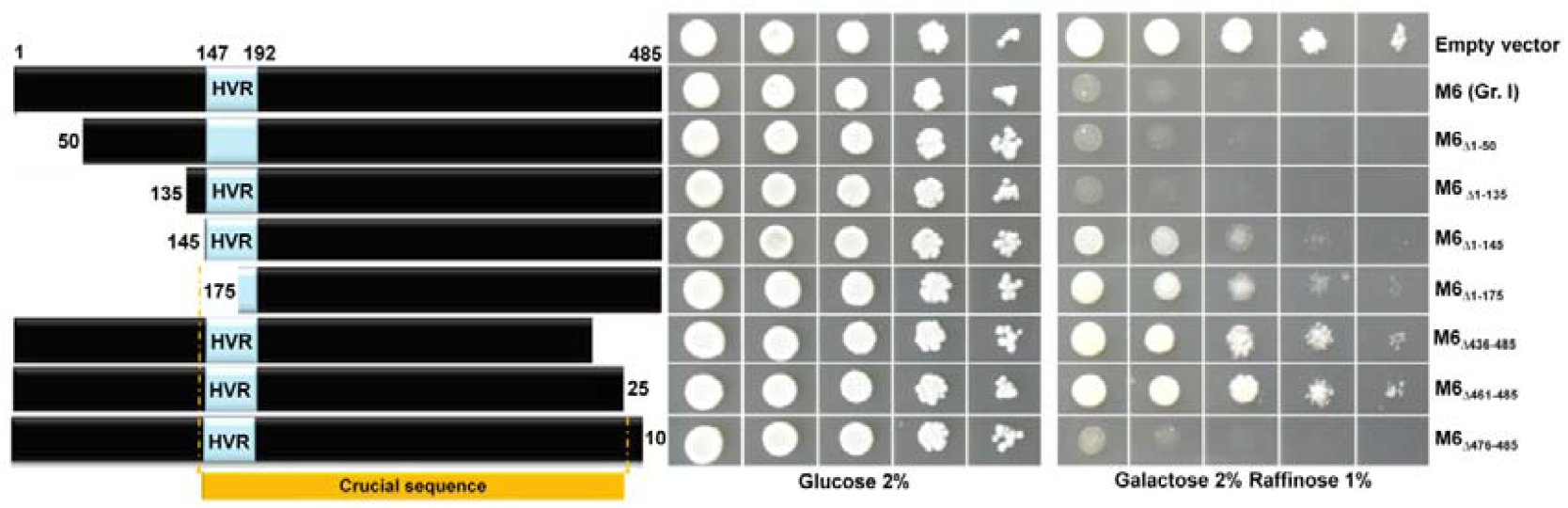
The HVR and the C-terminal region of AopW1 are required for cytotoxicity. Wild-type and shortened variants of *A. citrulli* M6 (group I) *aopW1* were cloned in pGML10 and transformed into *S. cereviasiae* BY4741 for growth inhibition assays. Yeast were grown in glucose (repressing) and in galactose/raffinose (inducing) medium. Pictures were taken after 3 days of growth at 28 °C and are representative of three independent experiments.

### Conserved residues in the HVR are crucial for induction of water-soaking-like cell death in *Nicotiana benthamiana*

Recently, Traore *et al*. (2019) showed that transient expression of group I and II AopW1 induces a water-soaking-like cell death phenotype in *N. benthamiana* leaves. Here we assessed the effects exerted by selected group I and II AopW1 variants carrying substitutions in some of the HVR a. a. positions (based on results from yeast inhibition assays). The AopW1 variants were agroinfiltrated into *N. benthamiana* leaves and their expression was verified by Western blot analysis (not shown).

In line with the results reported by Traore *et al*. (2019), both group I and II AopW1 induced water soaking in *N. benthamiana* leaves, with group I AopW1 inducing a much stronger effect (Figure 6). The group I variant of AopW1 carrying the three HVR substitutions that reduced cytotoxic activity in yeast [1 (V-154-I) + 2 (R-162-Q) + 3 (P-167-A)] did not differ considerably in terms of water soaking induction as compared with group I AopW1. However, the group II variant of AopW1 carrying the four substitutions that increased cytotoxicity of this effector in yeast [1 (I-154-V) + 2 (Q-162-R) + 3 (A-167-P) + 4 (T-174-A)] induced a stronger water-soaking phenotype in *N. benthamiana* leaves as compared with group II AopW1 (Figure 6). This result demonstrates the importance of the above a.a. positions for *in planta* cytotoxic activity of AopW1.

**Figure 6.**
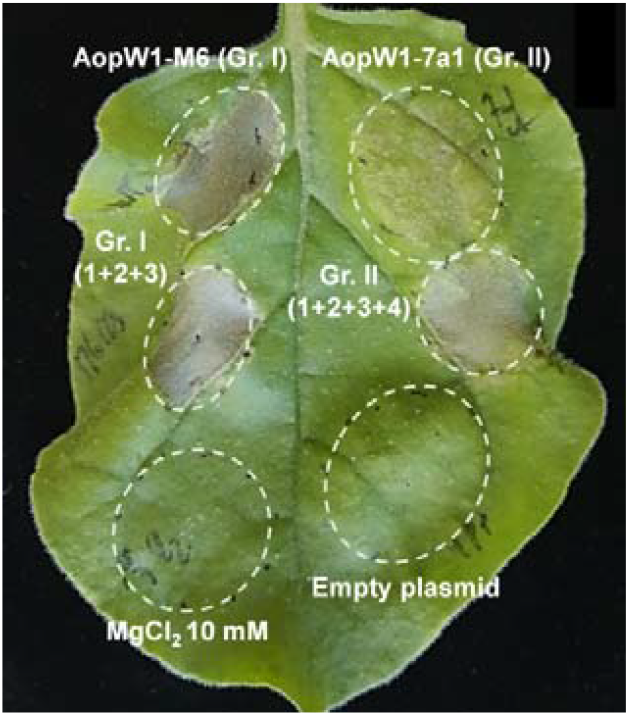
Transient expression of AopW1 and EHD1 in *N. benthamiana* leaves. **(A)** Wildtype or mutated variants of *A. citrulli* M6 (group I) and 7a1 (group II) AopW1 were transiently expressed in *N. benthamiana* leaves following agroinfection. See AopW1 variant details in Supplementary Table S2. Pictures were taken 3 days after infiltration.

### AopW1 localizes to the plant cell cytoplasm and interacts with chloroplasts

The subcellular localization of group I and II AopW1 was studied by confocal microscope analysis following agroinfiltration of *N. benthamiana* leaves. AopW1 ORFs were fused to yellow fluorescent protein (YFP) (Supplementary Table S2) for localization analyses. Overall, the work with group II AopW1 was easier than with group I AopW1, due to the extensive damage caused in *N. benthamiana* infiltrated sites by the latter, which challenged co-localization analysis.

Both group I and II AopW1 were located in the cytoplasm, with no nuclear localization being detected (Figure 7A and B). Interestingly, both effectors were shown to co-localize with chloroplasts. This was more clearly detected for group II AopW1 (Figure 7B), probably because of the aforementioned extensive damage caused by group I AopW1 to *N. benthamiana* cells. Co-localization of AopW1 with chloroplasts was in agreement with the detection of a chloroplast transit peptide (cTP) in the N-terminus of the effector, as detected by ChloroP 1.1 (Emanuelsson *et al*., 1999), WoLF PSORT (Horton *et al*., 2007) and Localizer (Sperschneider *et al*., 2017) softwares (Supplementary Appendix S1). Deletion of the first N-terminal 85 residues from group II AopW1, which contains the cTP sequence, abolished the co-localization of AopW1 with chloroplasts (Figure 7C).

**Figure 7.**
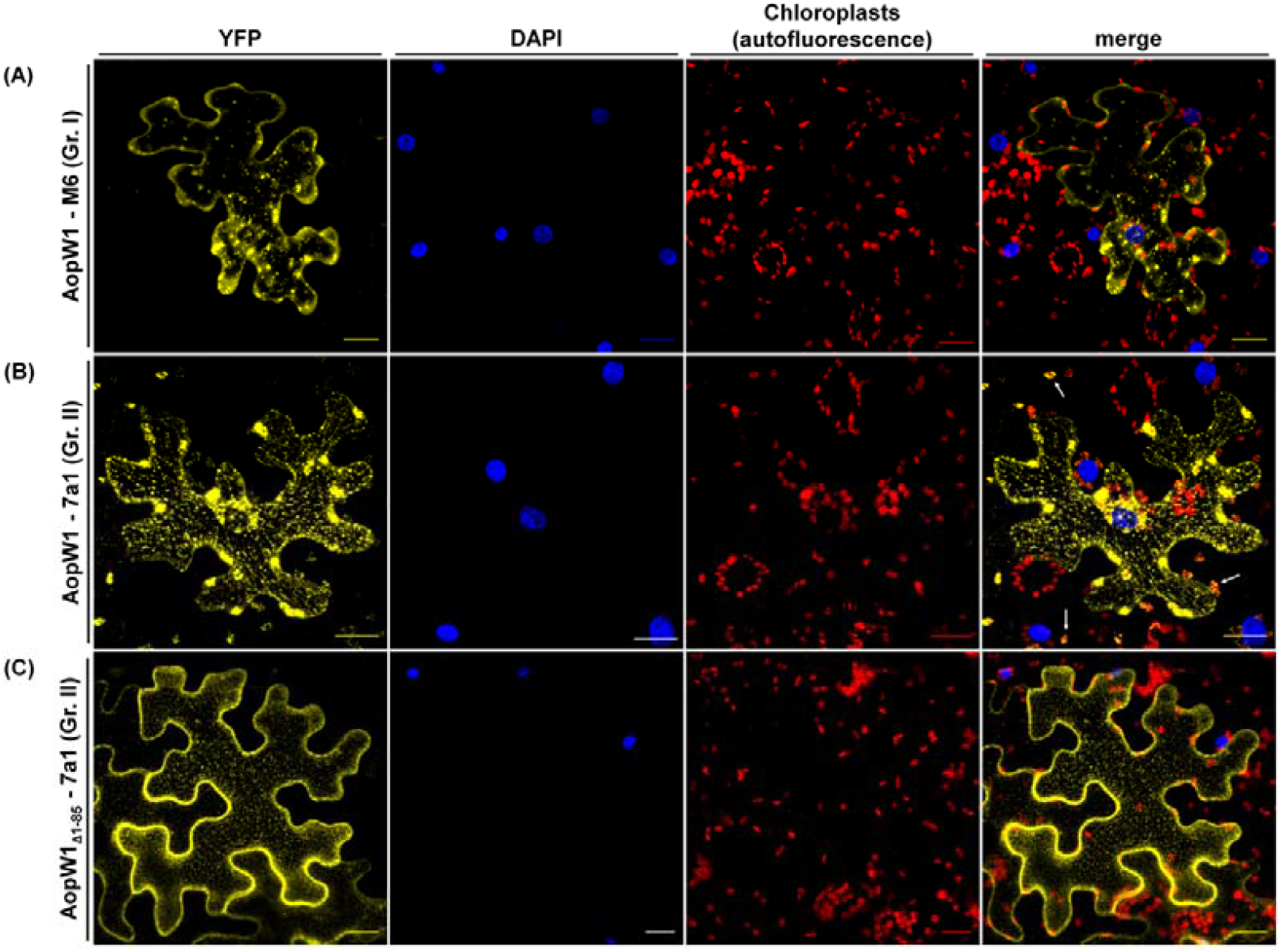
Subcellular localization of AopW1 in *N. benthamiana* leaf cells. Confocal images of *N. benthamiana* leaves 24 h after agroinfiltration. YFP-fused AopW1 from strains M6 (group I) and 7a1 (group II) are shown in yellow. Nuclei are shown in blue. Chloroplasts are shown in red. **(A)** Group I AopW1. (**B**) Group II AopW1. (**C**) A short variant of group II AopW1 lacking the predicted chloroplast transit peptide (cTP) in its N-terminus (AopW1_Δ1-85_). Removal of this region causes a delocalization of AopW1 into chloroplasts. White arrows indicate colocalization of AopW1 with chloroplasts. Scale bars indicate 20 μm.

### Group I AopW1 disrupts the plant actin cytoskeleton and affects endoplasmic reticulum organization

To evaluate the interaction of AopW1 with the plant actin cytoskeleton, we coexpressed AopW1 fused with YFP with the plant actin marker, DsRed-ABD2 (Voigt *et al*., 2005a) following agroinfiltration of *N. benthamiana* leaves. While both group I and II AopW1 co-localized with the actin marker, only group I AopW1 disrupted the actin cytoskeleton *in planta* (Figure 8A).

**Figure 8.**
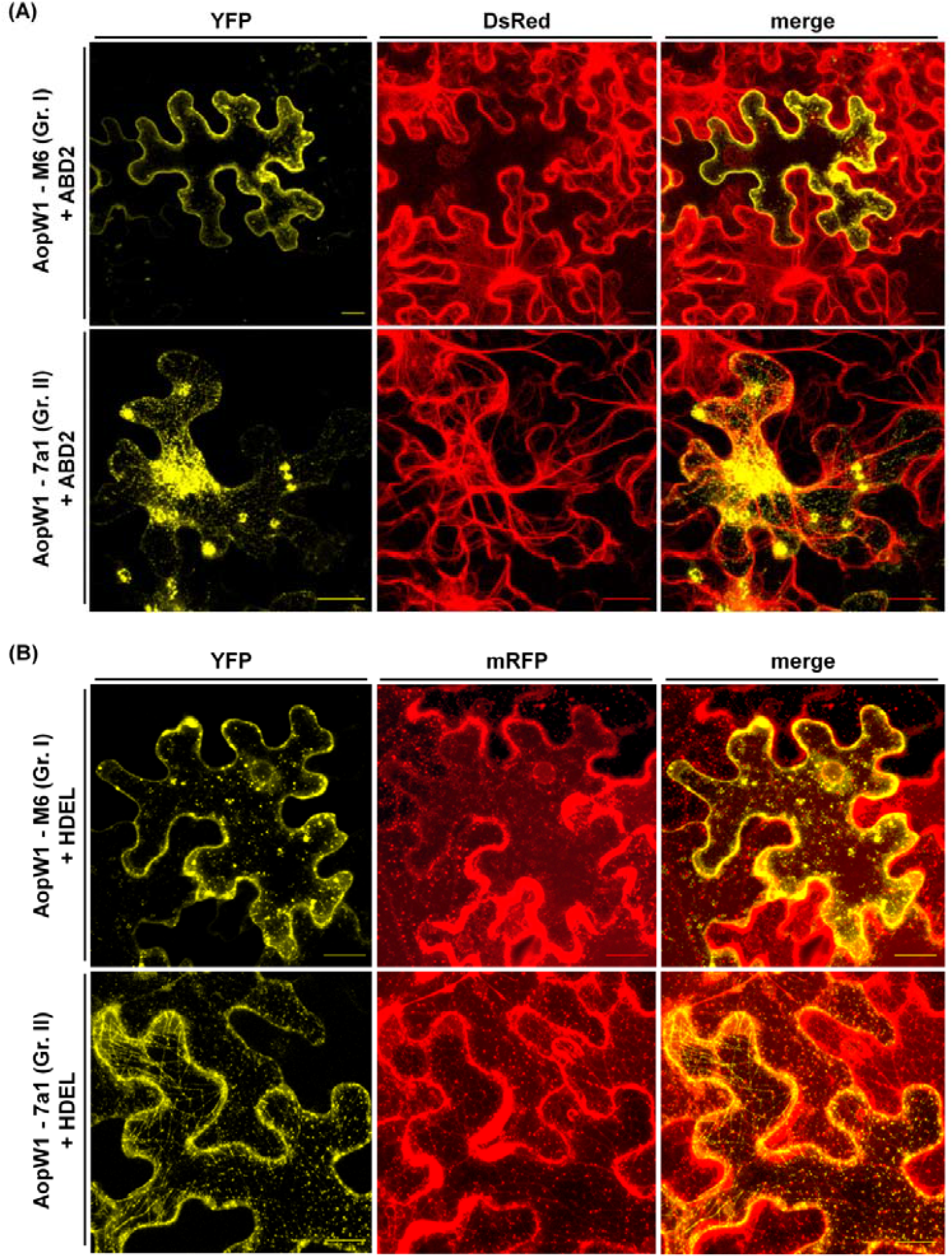
Group I AopW1 disrupts the actin cytoskeleton and alter the endoplasmic reticulum (ER) distribution in *N. benthamiana* leaf cells. Confocal images of *N. benthamiana* leaves 48 and 24 h after agroinfiltration with plant markers and AopW1, respectively. YFP-fused AopW1 from strains M6 (group I) and 7a1 (group II) are shown in yellow. The actin marker DsRed-ABD2 (**A**) and the ER marker mRFP-HDEL (**B**) are shown in red. Scale bars indicate 20 μm.

Proteins involved in actin polymerization are often associated with the endoplasmic reticulum (ER) (Zhang *et al*., 2013). To assess whether AopW1 interacts with the ER, we carried out co-expression assays of AopW1-YFP with the ER marker mRFP-HDEL (Runions *et al*., 2006; Schoberer *et al*., 2009). While both versions of AopW1 colocalized with the ER, only group I AopW1 caused a disorganization of the usual ER architecture (Figure 8B).

### AopW1 co-localizes with plant endosomes

The plant actin cytoskeleton plays an essential role in several biological processes in plants, including endocytosis and endosomal trafficking (Paez-Garcia *et al*., 2018; Thomas *et al*., 2009). We tested co-localization of AopW1-YFP with various plant endosome markers fused with several fluorescent proteins in *N. benthamiana* leaf cells. The tested markers were FYVE-DsRed, which binds phosphatidyl inositol (Voigt *et al*., 2005a); AtEHD1-CFP, which partially co-localizes with FYVE, RabD2b (Wave33) and RabA1e (Wave34), and functions in endocytic recycling (Bar *et al*., 2008a,b; Bar *et al*., 2013); mCherry-Wave33, which possesses endosomal and trans-Golgi localization (Bar *et al*., 2013); and Ara7 (RabF2b)-CFP, which likely localizes to recycling and late endosomes (Bar *et al*., 2013; Lee *et al*., 2004).

Expression of group I AopW1 in *N. benthamiana* leaves revealed this effector partially co-localizes with all tested endosome markers, suggesting AopW1 possibly interferes with early and recycling endosomes (Figure 9A). In contrast, we were not able to detect co-localization of group II AopW1 with DsRed-FYVE (Figure 9B). The pattern of distribution of group II AopW1 appeared to be similar to those of markers mCherry-Wave33 or Ara7-CFP, which could be due to their actin-dependent localization. Nevertheless, unlike the observed for group I AopW1, no clear colocalization signals were detected between these markers and group II AopW1 (Figure 9).

**Figure 9.**
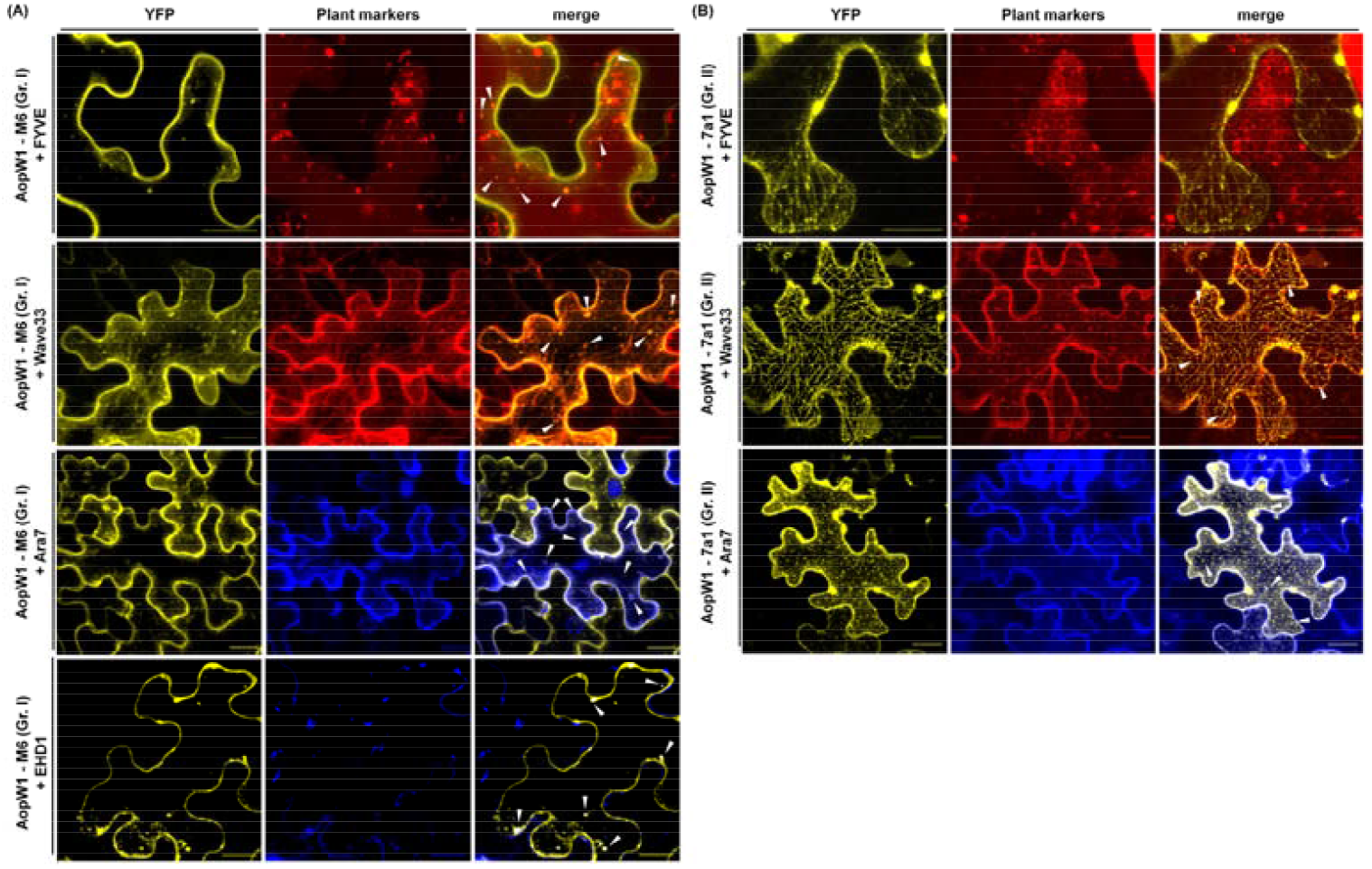
AopW1 co-localizes with plant endosomes in *N. benthamiana* leaf cells. Confocal images of *N. benthamiana* leaves 48 and 24 h after agroinfiltration with plant markers and AopW1, respectively. YFP-fused AopW1 from strains M6 (group I, **A**) and 7a1 (group II, **B**) are shown in yellow. Endosome markers DsRed-FYVE and mCherry-Wave33 (RabD2b) are shown in red. Endosome markers Ara7-CFP (RabF2b) and AtEHD1-CFP are shown in blue. White arrows indicate co-localization of AopW1 with plant endosomes. Scale bars indicate 20 μm.

### EHD1 attenuates AopW1 cytotoxicity and increases immune responses

Observation of leaves used in co-localization experiments suggested that coinfiltration of group I AopW1 with the AtEHD1-CFP marker attenuated the toxic effect induced by the effector. To verify these findings, we carried out additional leaf infiltration assays in which *A. tumefaciens* carrying group I or II AopW1 were infiltrated alone or in combination with *A. tumefaciens* carrying AtEHD1. Coinfiltrations were carried out at the same site or at a close distance between bacteria carrying the two proteins. Results from these experiments confirmed that expression of EHD1 in *N. benthamiana* attenuated the toxic effect induced by group I AopW1, and completely abolished the toxicity induced by group II AopW1 (Figures 10A and B).

**Figure 10.**
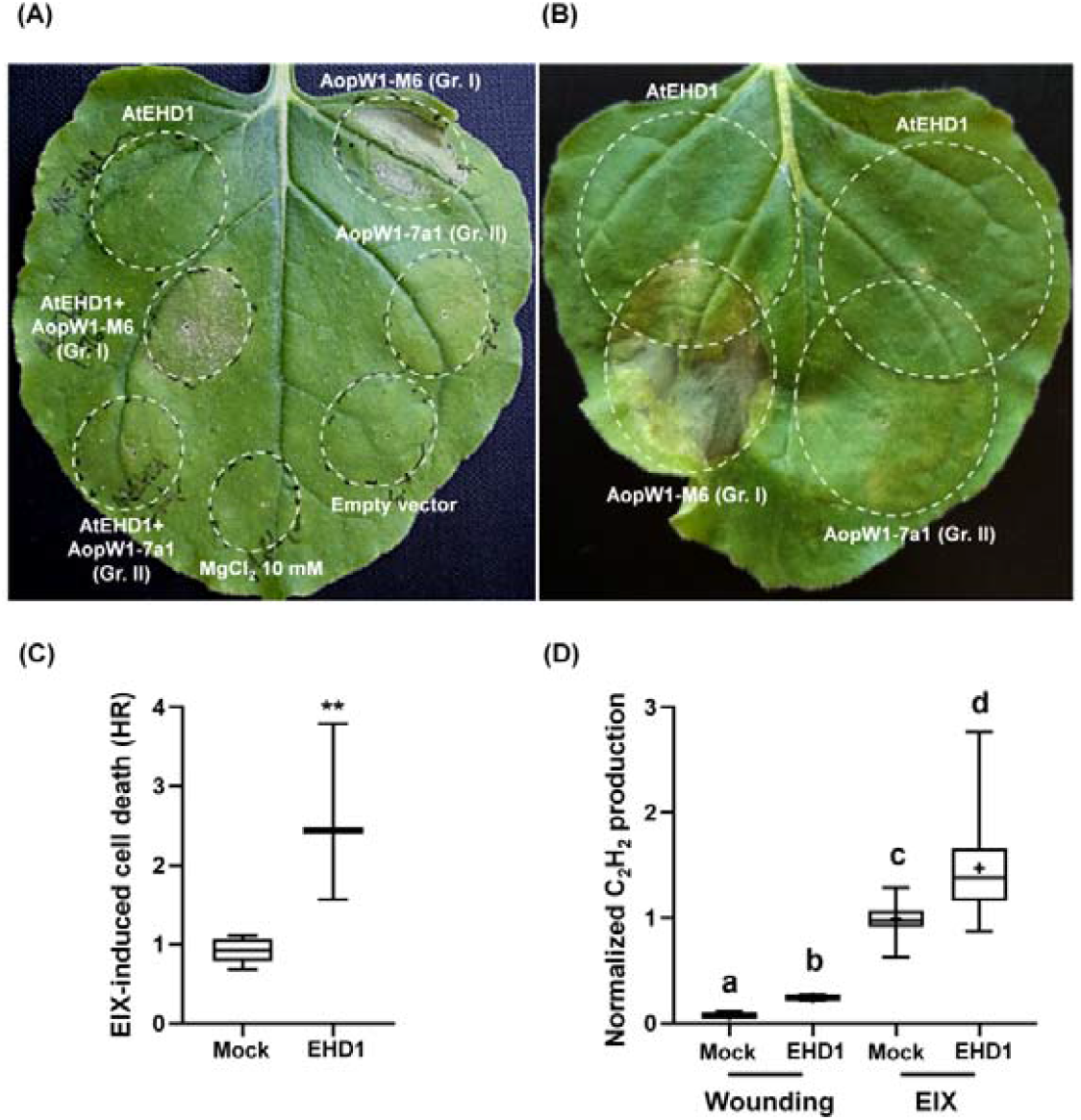
EHD1 attenuates AopW1 cytotoxicity and increases defence responses. Transient expression of EHD1 in *N. benthamiana* attenuates or abolishes the water soaking phenotype triggered by group I and group II AopW1, respectively. *Agrobacterium*-mediated transient expression of EHD1 and AopW1 from *A. citrulli* M6 (group I) and 7a1 (group II) by infiltration of individually **(A)** or overlapping **(B)** *A. tumefaciens* cultures. Pictures were taken 3 days after infiltration. Transient expression of EHD1 in *N. tabacum* leaves increases EIX-induced cell death (hypersensitive response, HR) **(C)** and EIX-induced ethylene biosynthesis **(D)**. For HR assays, *N. tabacum* leaves were transiently transformed to co-express EIX and EHD1-GFP or free GFP (mock). HR development was quantified 72 h post-injection. Mock average HR level was defined as 1. Data are means and SE of 3 independent experiments, with asterisks denoting significant difference from mock in a two-tailed t-test (N=3, **p=0.0062). For ethylene biosynthesis assays, leaf disks of transiently transformed *N. tabacum* leaves with EHD1 or mock (free GFP) were floated on a solution with 250 mM sorbitol without (wounding) or with 1 μg/mL EIX, 48 h post transformation. Ethylene biosynthesis was measured after 4 h. The mock + EIX average induction value was defined as 1. Data represent means and SE of 3 independent experiments, with letters denoting significant differences among treatments in a one-way ANOVA with a Dunnett post-hoc test (N=21, p<0.0001). Box-plots display minimum to maximum values, with inner quartile ranges indicated by box and outer-quartile ranges by whiskers. Line indicates median, “+” indicates mean.

EHD1 is known to affect endosomal recycling in *Arabidopsis* (Bar *et al*., 2008a; Bar *et al*., 2013). We asked whether EHD1 could be directly involved in plant immunity. For this purpose, we transiently co-expressed EHD1 and the fungal defence elicitor ethylene-inducing xylanase (EIX; Dean *et al*., 1991; Ron and Avni, 2004) in leaves of *Nicotiana tabacum*, and monitored EIX-induced cell death 72 h post-transformation. Leaves co-expressing EIX and EHD1 showed significantly higher levels (p=0.0062) of induced cell death than leaves expressing EIX alone (Figure 10C). Similarly, *N. tabacum* leaves transiently transformed with EHD1 showed significantly increased levels (p<0.0001) of ethylene biosynthesis than non-transformed leaves following treatment with EIX (Figure 10D).

### Group I AopW1 reduces callose deposition in *N. benthamiana* leaves after PTI induction

Increased callose deposition is a common marker of PTI and many pathogen effectors are able to reduce callose deposition upon induction of PTI (Voigt, 2014; Wang *et al*., 2021). To assess whether AopW1 affects callose deposition, *N. benthamiana* leaves were treated with the flagellin-derived PTI elicitor flg22 (Felix *et al*., 1999). After 24 h, leaves were agroinfiltrated with group I or II AopW1. Leaf samples were collected after 24 h for assessment of callose deposition. As controls, leaves were left untreated or infiltrated with *A. tumefaciens* GV3101 carrying pEarleyGate 101 (empty vector). Group I AopW1 significantly (p < 0.05) reduced the number of callose deposits induced by flg22, as compared with controls. In contrast, group II AopW1 did not significantly affect callose deposition as compared with controls (Figure 11).

**Figure 11.**
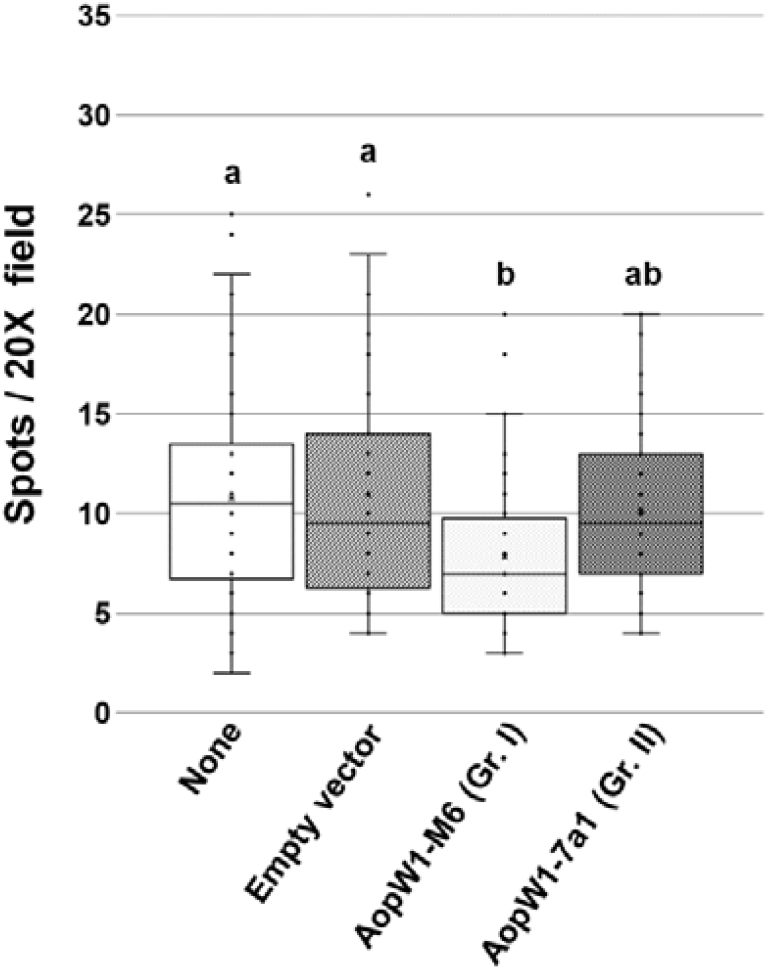
Group I AopW1 reduces callose deposition in *N. benthamina* cells after PTI induction. *Nicotiana benthamiana* leaves were treated with 40 μM flg22 for 24 h and then coinfiltrated with *A. tumefaciens* GV3101 carrying pEarleyGate101, either empty or containing the *aopW1* ORFs of *A. citrulli* M6 (group I) or 7a1 (group II). After 24 h post agroinfiltration, 1 cm-diameter disks were collected, stained with aniline blue and callose deposits were quantified. Callose deposits were counted from 6 areas/3 leaves/3 plants. Data represent means and SE from one experiment out of two with similar results. Letters indicate significant (p=0.05) differences among treatments by ANOVA, using both Bonferroni and HSD Tukey tests.

## DISCUSSION

*Acidovorax citrulli* requires a functional T3SS to cause disease (Bahar and Burdman, 2010; Johnson *et al*., 2011). A large number of novel T3E genes have been recently identified in the genome of the *A. citrulli* model strain M6 (Jiménez-Guerrero *et al*., 2020). M6 belongs to one of the two predominant groups of *A. citrulli* strains, group I, which is composed of strains that were mainly isolated from melon but also from other non-watermelon cucurbits. The second major group of strains, group II, is composed of strains that show preferential association with watermelon, and can be clearly distinguished by genetic and biochemical traits from group I strains (Burdman *et al*., 2005; Eckshtain-Levi *et al*., 2016; Feng *et al*., 2009; Walcott *et al*., 2000; Walcott *et al*., 2004). We have shown that group I and II strains differ in their T3E arsenal. Firstly, several T3Es appear to be group-specific. Secondly, many T3E genes that are shared by the two groups show significant differences in their sequences (Eckshtain-Levi *et al*., 2014; Jiménez-Guerrero *et al*., 2020). In this study, we characterized an *A. citrulli* T3E, AopW1, which belongs to the second category.

AopW1 shares similarity with *P. syringae* HopW1, and with T3Es from other plant-pathogenic bacterial species, including *Acidovorax* spp. (Jiménez-Guerrero *et al*., 2020; Figure 3). This effector was recently shown to contribute to *A. citrulli* virulence (Traore *et al*., 2019). In the present study, heterologous expression in yeast revealed that while group I AopW1 is extremely toxic to these cells, the group II version of this effector has only a minor toxic effect (Figure 1).

Among AopW1 homologous effectors, the only one that was well characterized is *P. syringae* HopW1. While *aopW1* encodes a product of 485 amino acids (a.a.), HopW1 is 774 a.a-long. In fact, AopW1 shares similarity with the central and C-terminal parts of HopW1. Lee *et al*. (2008) showed that HopW1 triggers immunity responses in the *Arabidopsis* Ws ecotype and in *N. benthamiana*. The authors also showed that HopW1 interacts with a putative acetylornithine transaminase (WIN1), a protein phosphatase (WIN2) and a firefly luciferase superfamily protein (WIN3) of *Arabidopsis*, with the HopW1 C-terminal domain being required for these interactions. Later, Kang *et al*. (2014) showed that HopW1 promotes virulence in a different *Arabidopsis* ecotype, Col-0, by inhibiting actin polymerization and severely disrupting endosome trafficking.

AopW1 is highly conserved among group I and II strains of *A. citrulli*, except for a hypervariable region (HVR) located at a.a. positions 147-192, and showing 14 a.a. differences between group I and II strains (Figure 3B). Importantly, the HVR is included in the part of AopW1 that shares similarity with the central/C-terminal regions of HopW1 that are required for cytotoxic activity. Here we showed that deletion of the HVR abolishes the ability of group I AopW1 to exert toxic effects on yeast (Figure 5). We also noticed that among the 14 variable a.a. positions between group I and II AopW1 in the HVR, six are well conserved between group I AopW1, HopW1 and other HopW1 homologs (Figure 3B). We therefore hypothesized that these positions are critical for the activity of AopW1 and other effectors of this family. In support of this notion, growth inhibition assays of yeast expressing mutated versions of AopW1 revealed that a combination of three substitutions in the HVR (V-154-I, R-162-Q and P-167-A) were sufficient to cause a significant reduction of cytotoxicity exerted by group I AopW1. On the other hand, combination of four substitutions (I-154-V, Q-162-R and A-167-P and T-174-A) conferred high cytotoxic ability to the group II version of AopW1 (Figure 4). These findings demonstrate the important role of the above a. a. positions for the cytotoxic activity of effectors belonging to the HopW1 family.

The actin cytoskeleton is a complex network of dynamic polymers that play important roles in a wide range of cellular processes (Mishra *et al*., 2014; Pollard and Borisy, 2003). The actin cytoskeleton also plays an essential role in plant immunity, assisting in multiple defence functions in both early and late defence responses, such as vesicle trafficking and endo/exocytosis, fortification of the cell wall and deposition of callose (Hardham *et al*., 2007; Li and Staiger, 2018). Upon bacterial infection, epidermal cells of *Arabidopsis* leaves show early increase in density of actin filaments and late actin remodelling, with these responses being correlated with PTI and effector-triggered susceptibility, respectively (Henty-Ridilla *et al*., 2013). Therefore, it is not surprising that pathogen effectors target host cytoskeletal organization in order to subvert plant defence responses, as demonstrated for HopW1 (Kang *et al*., 2014) but also for a different *P. syringae* effector, HopG1 (Shimono *et al*., 2016). In addition, several studies indicate that other *P. syringae* effectors, HopAV1, HopAZ1, HopZ1a and HopE1, as well as AvrBsT from *Xanthomonas euvesicatoria*, or the powdery mildew effector ROPIP1, could be acting to some extent on the plant cell cytoskeleton (Cheong *et al*., 2014; Choi *et al*., 2017; Guo *et al*., 2016; Lee *et al*, 2012; Nottensteiner *et al*., 2018). Accordingly, we showed that group I AopW1 disrupts muscle F-actin *in vitro* (Figure 2), and affects yeast and plant filamentous actin (F-actin) *in vivo* (Figures 8A and 11; Supplementary Figure S1). Findings from our study also support the notion that group II AopW1 is an attenuated version of this effector in terms of actin disruption ability (Figures 2 and 8A; Supplementary Figure S1).

One of the plant responses to pathogen invasion is fortification of the cell wall through callose deposition (Voigt, 2014). This immune response is assisted by the actin cytoskeleton (Li and Staiger, 2018). Pathogens utilize protein effectors to suppress plant immune response, including reduction of callose deposition (Wang *et al*., 2021). Here we showed that group I but group II AopW1 is able to significantly reduce callose deposition in *N. benthamiana* leaves pre-treated with the PTI elicitor flg22 (Figure 11). This finding could be explained, at least partially, by the severe disruption of the actin cytoskeleton by group I but not by group II AopW1. Differences between the two versions of the effector were also observed in alteration of the ER structure, which could be indirectly associated with the differences between the effectors in actin disruption ability (Figure 8B).

Actin is directly involved in vesicle trafficking and endo/exocytosis processes that are essential for the delivery of specialized proteins and defence molecules to the cytoplasmic membrane (Li and Staiger, 2018). Several defence receptors are internalized into early endosomes, and further recycled back to the membrane via recycling endosomes, or targeted for degradation via the late endosome pathway. Increase of the amount of a receptor in endosomes causes a concomitant signalling enhancement, whereas abolishment of endosome formation once the receptor is internalized causes signalling attenuation (Bar and Avni, 2014). Therefore, endosome components appear to be attractive targets of pathogen effectors. As mentioned above, HopW1 was shown to reduce the number of endosome vesicles *in planta* (Kang *et al*., 2014). Other T3Es seem to be associated with plant endosomes, such as HopM1 from *P. syringae*, which interferes with AtMIN7/BEN1 function at the early endosome/trans-Golgi network (TGN) (Nomura *et al*. 2006; Nomura *et al*. 2011). Here we showed that group I AopW1 clearly co-localizes with the endosome markers FYVE, Ara7, Wave33 and EHD1, supporting that it acts in some way in early and recycling endosome functioning (Figures 9 and 12).

**Figure 12.**
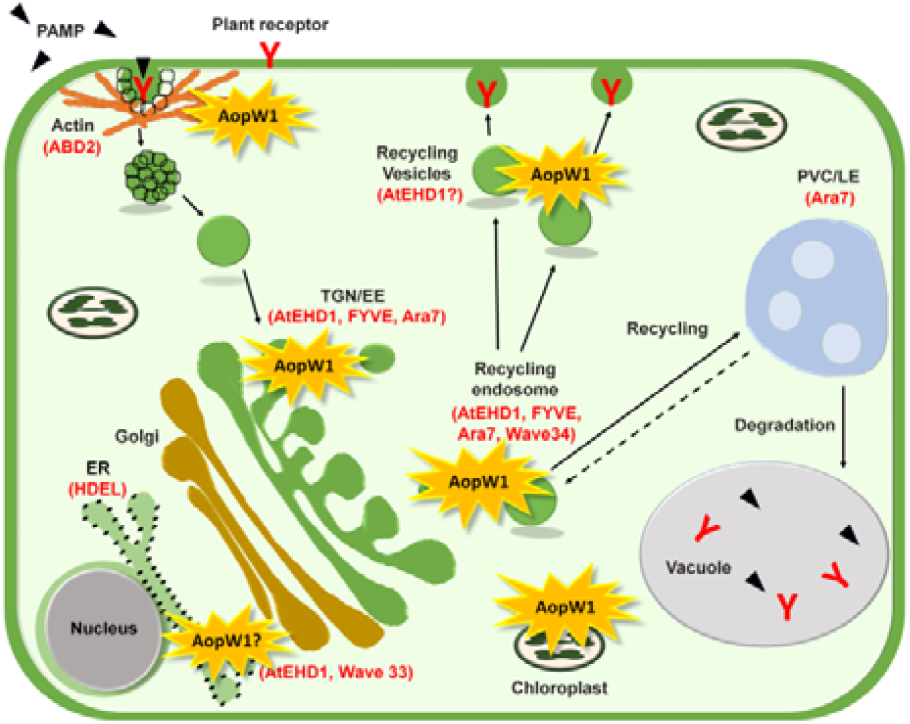
Schematic representation of AopW1 localization and targeting in the plant cell. AopW1 localizes at the cell cytoplasm, where it disrupts actin filaments and co-localizes to early and recycling endosomes. AopW1 also localizes to chloroplasts. Co-localization with known markers is indicated in red (AtEHD1, FYVE, Ara7 and Wave33). Arrows point to trafficking pathways from the membrane to the early/re-cycling endosome and back to the membrane. TGN/EE, trans-Golgi network/early endosome; MVBs/LE, multivesicular bodies/late endosome.

Co-localization of EHD1with FYVE and Wave33 was reported by Bar *et al*. (2008; 2013), who suggested a role for this protein in endocytic recycling. The mammalian EHD1 homologs are known to be involved in endocytic recycling (Galperin *et al*., 2002; Grant and Caplan, 2008). Moreover, it has been shown that overexpression of EHD1 protects against salinity stress, suggesting an association between endocytic recycling and plant stress coping mechanisms (Bar *et al*., 2013). Here we showed that overexpression of EHD1 attenuates the cytotoxic effect of group I AopW1 and abolishes the cytotoxic effect of group II AopW1 in *N. benthamiana* leaves (Figures 10A and B), thus supporting the involvement of this effector in endocytic recycling. In addition to the recycling mechanism which may underlie the attenuated effector cytotoxicity, we show that transient expression of EHD1 in *N. tabacum* leaves increases defence responses induced by the EIX elicitor (Figures 10C and D), suggesting that increased endocytic recycling may increase plant immunity. Overall, our study provides first evidence on the potential role of the endosome-associated protein EHD1 in regulating immune responses. The attenuating effect of AopW1 exerted by EHD1 could be mediated by different mechanisms such as toxicity relief through endocytic recycling and/or induction of host immunity.

Chloroplasts play an important role in plant immunity as they are the source of important defence signalling molecules (Nomura *et al*., 2012; Serrano *et al*., 2016; Singh *et al*., 2018). AopW1 possesses a predicted cTP signal in its N-terminal part (Supplementary Appendix 1) and co-localization of AopW1 to chloroplasts in a cTP signal-dependent manner was demonstrated in this study (Figures 7C). Whether AopW1 interferes with chloroplast functioning or with a chloroplast signalling pathway is yet to be determined.

In conclusion, our study provides insights into mechanistic features of T3Es belonging to the HopW1 family. While it was previously shown that group I and II strains of *A. citrulli* possess different versions of AopW1 (Eckshtain-Levi *et al*., 2014), here we show that these versions substantially differ in their ability to interact with plant cell compartments and disrupt the actin cytoskeleton (Figure 12). It is yet to be elucidated whether the differences between the two versions of AopW1 are important determinants for the distinguished patterns of host preferential association of the two groups of *A. citrulli* strains. Considering the subtle contribution of most bacterial effectors (including AopW1) to virulence, and the high amount of T3Es in *A. citrulli*, this is a very challenging task. The question also remains whether the variation in AopW1 is among the significant evolutionary alterations that occurred during the process of adaptation of *A. citrulli* to different hosts. In this regard, it is worth remarking that a similar variation in the AopW1 HVR could be detected among strains of *A. avenae*, which are able to infect several graminaceous plants.

## EXPERIMENTAL PROCEDURES

### Strains, plant material and growth conditions

Strains and plasmids used in this study are listed in Supplementary Table S2. *Escherichia coli* and *Agrobacterium tumefaciens* as well as derived strains were cultured in Luria-Bertani (LB; Difco Laboratories, USA) medium at 37 °C and 28 °C, respectively. *Acidovorax citrulli* strains were grown in nutrient broth (NB; Difco Laboratories) or nutrient agar (NA; NB containing 15 g/l agar) at 28 °C. When required, the media were supplemented with the antibiotics ampicillin (Ap, 100 μg ml^-1^ for *E. coli*, and 200 μg ml^-1^ for *A. citrulli*), rifampicin (Rif, 50 μg ml^-1^), kanamycin (Km, 30 μg ml^-1^ for *E. coli* and 50 μg ml^-1^ for *A. tumefaciens*), and gentamycin (Gm, 10 μg ml^-1^). *Saccharomyces cerevisiae* BY4741 (Brachmann *et al*., 1998) was grown at 30 °C on YPD medium (Rédei, 2008). For repressing and inducing conditions, BY4741 derivate strains were grown in selective synthetic complete medium without leucine supplemented with 2% glucose or 2% galactose and 1% raffinose, respectively (Salomon and Sessa, 2010). *Nicotiana benthamiana* plants (Goodin *et al*., 2008) were grown in a growth chamber under the following controlled conditions: 16 □h at 26□°C in the light and 8□h at 18□°C in the dark, 70% humidity. *Nicotiana tabacum* cv. Samsun NN plants were grown from seeds under greenhouse long day conditions (16 h light/ 8 h dark), at 25 □°C.

### Molecular manipulations

Routine molecular manipulations and cloning procedures were carried out by standard techniques unless stated otherwise. Restriction enzymes and T4 DNA ligase were purchased from Fermentas (Thermo Fisher Scientific, USA). AccuPrep^®^ Plasmid Mini Extraction Kit (Bioneer Corporation, Republic of Korea) and Wizard^®^ SV Gel and PCR Clean-Up System (Promega Corporation, USA) were used for plasmid and PCR product extraction and purification, respectively. Bacterial DNA was extracted using the GeneElute Bacterial Genomic DNA Kit (Sigma-Aldrich, USA). All constructs were verified by DNA sequencing at Hy Laboratories (Israel). PCR primers were purchased from Hy Laboratories or Sigma-Aldrich Israel. All oligonucleotides primers used in this study are listed in Supplementary Table S4. PCR reactions were performed with the Readymix Red Taq PCR reactive mix (Sigma-Aldrich), Phusion high-fidelity DNA polymerase (Fermentas) or with the Q5^®^ High-Fidelity DNA Polymerase (New England Biolabs, USA) using an Eppendorf (Germany) thermal cycler.

For immunostaining, proteins were separated by SDS-PAGE and electroblotted using the iBlot Gel Transfer Stacks (Invitrogen, USA) and iBlot Transfer Device (Invitrogen), following manufacturer’s instructions. Nitrocellulose membranes were blocked with TBS containing 0.1 % (v/v) Tween 20 and 3% (w/v) skim milk and incubated with antibodies raised against c-Myc, His and HA (Cell Signalling Technology, USA) diluted 1:1000 in the same solution. Anti-mouse or anti-rabbit HRP-linked antibodies (Cell Signalling Technology) were used as secondary antibodies. Reactions were visualized using a chemiluminescent substrate (Cyanagen, Italy) in a LAS500 apparatus (GE Healthcare, USA).

### Cloning and transformation of yeast and bacteria

For expression in yeast, the *aopW1* ORFs of *A. citrulli* M6 and 7a1 were PCR-amplified with specific primers. The obtained products were inserted into the *Bam*HI/*Eco*RI sites of pGML10 (Iha and Tsurugi, 1998), following pretreatment of the inserts and plasmids with the same restriction enzymes. The ORFs of the *hopW1* gene of *P. syringae* pv. *maculicola* ES4326 and of the *hopW1* homologous gene of *X. translucens* pv. *translucens* (*Xtt*) DSM 18974 were cloned into pGML10 by restriction free (RF) method (van den Ent and Lowe, 2006) using appropriate primers, and plasmid pBAV154-dexHopW1-HA or genomic DNA of *Xtt* DSM 18974 as templates, respectively. The mutated variants of AopW1 were generated by inserting mutations in the corresponding primers in combination with the Quick Change Lightening Site Directed Mutagenesis kit (Agilent Technologies, SC, USA). For this purpose, PCR reactions for each mutation were carried out in 50 μl reaction mixture containing 5 μl of 10X reaction buffer, 100 ng of template DNA, 125 ng of both oligonucleotide primers, 1 μl of dNTP mix, 1.5 μl of Quickchange Solution reagent and 1 μl of Quick Change lightening enzyme. The final volume was made with sterilized distilled water (SDW). The cycling parameters included an initial denaturation step at 95 °C for 2 min, 18 cycles of denaturation (at 95 °C for 20 s), annealing (at 60 °C for 10 s) and extension (at 68 °C for 2.5 min), and final extension at 68 °C for 5 min. Then, the amplified plasmid DNA product was digested with *Dpn*I provided with the kit. Effector gene variants carrying combinations of individual mutations in the background of the M6 and 7a1 *aopW1* genes were generated using the same procedure. The resulting plasmids were mobilized into *S. cerevisiae* BY4741 by the lithium acetate yeast transformation method (Gietz *et al*., 1992). Transformed yeast cells were plated onto selective synthetic complete without leucine medium supplemented with 2% glucose.

For *Agrobacterium*-mediated transient expression, the *aopW1* ORFs of strains M6 and 7a1 were amplified with appropriate primers. The resulting products were cloned into pDONR207 entry vector and then mobilized into pEarlyGate 101 binary vector (Earley *et al*., 2006) by the Gateway cloning system (Thermo Fisher Scientific). The generated plasmids were then transformed into *A. tumefaciens* GV3101 by heat shock transformation as described (Zhou *et al*., 2009).

For expression and purification of recombinant AopW1 to assess F-actin *in vitro* disruption activity, the *aopW1* ORFs without their 300-bp N-terminal region were amplified by PCR with specific primers. The resulting fragments were cloned into the *Eco*RI/*Xho*I site of pET28a, leading to generation of recombinant AopW1 fused to polyhistidine tag (His-tag) in both extremes. The generated plasmids were mobilized into *E. coli* BL21(DE3) (Studier and Moffatt, 1986) by the heat shock method (Froger and Hall, 2007).

### Yeast growth inhibition and viability assay

Growth inhibition assays were performed as described (Salomon *et al*., 2011). *Saccharomyces cerevisiae* BY4741 carrying pGML10 with *aopW1* ORFs were grown at 30 °C overnight in liquid selective medium containing 2 % glucose. Cultures were centrifuged (800 *g*, 5 min, at room temperature; twice) and pellets were suspended with SDW to an optical density at 600 nm (OD_600_) of 1.0. For each culture, four serial dilutions were prepared and 10-μl aliquots from each dilution were spotted onto solid selective medium containing 2% glucose (repressing medium), or 2% galactose and 1% raffinose (inducing medium), or onto inducing medium supplemented with 7 mM caffeine, 1 M sorbitol or 0.5 M sodium chloride. Yeast cells were incubated at 30 °C for 3 days.

For yeast viability assays, overnight cultures of yeast (as detailed above for growth inhibition assays) were diluted with fresh repressing medium to an OD600 of 0.5, and incubated for 2 h at 30 °C with shaking (200 rpm). Yeast cells were washed as described above but pellets were resuspended in 2 ml of induction medium to an OD_600_ of 0.2. The resulting cultures were incubated at 30 °C with shaking (200 rpm) and 0.1-ml aliquots were collected at different time points. The concentrations of viable cells were determined by plating of serial dilutions on selective repressing medium, with plates being incubated for 2 days at 30 °C. Data was analysed by Student t-test. The expression of the effectors was validated by Western blot analysis, following the procedure described by Salomon and Sessa (2010), using a c-Myc primary antibody (not shown).

### Yeast actin staining

Yeast actin staining was performed as described (Adams and Pringle, 1991), with some modifications. Briefly, yeast overnight cultures grown as described above were diluted with fresh repressing medium to and OD_600_ of 0.4 and incubated for 2 h at 30 °C with shaking (200 rpm). Yeast cells were washed as described above and pellets were resuspended in 4 ml inducing medium to an OD_600_ of 0.2. Cultures were incubated at the same conditions for 8 h to allow effector expression. During this period, cultures were refreshed with inducing medium at least once. Cultures were then centrifuged (800 *g*, 5 min, at room temperature), and pellets were washed with phosphate buffered saline solution (PBS; 137 mM NaCl, 2.7 mM KCI, 8 mM Na_2_HPO_4_ and 2 mM KH_2_PO_4_; pH 7.2). Pellets were carefully resuspended in 1 ml of fresh paraformaldehyde 4% solution in PBS. Fixation was done by incubation at 4 °C for 15 min with soft rotation. Cells were washed by centrifugation (500 *g*, 5 min, at room temperature) and resuspension with PBS twice, and finally resuspended in 1 ml of fresh PBS with 0.1% Triton X-100. The suspensions were incubated at room temperature for 20 min, after which the cells were washed as described above with PBS (twice). For staining of yeast cell actin and nucleus, the pellets were resuspended with 100 μl of blocking solution (1X PBS, 1% BSA and 0.1% saponin) containing 50 μg/ml of TRITC-phalloidin (Sigma-Aldrich) and 2.5 μg/ml of 4’,6-diamidino-2-phenylindole (DAPI, Sigma-Aldrich) (Kapuscinski and Skoczylas, 1977). The suspensions were incubated for 1 h at 4 °C, at dark conditions. Cells were then washed twice by centrifugation and resuspension with PBS as described above, and finally resuspended in 5-10 μl of 0.1 M propyl gallate in 100% glycerol. One-microliter drops of the suspensions were visualized in a Leica SPE Confocal Microscope (Leica Microsystems, Germany), using poly-L-lysine microscope slides.

### Expression and purification of recombinant AopW1

Expression and purification of recombinant His-tagged AopW1 was performed following the protocol described by the Macherey Nagel (Germany) purification of His-tag protein manual. For expression of AopW1, *Escherichia coli* BL21(DE3) carrying plasmids pET28a::_M6_*aopW1-M6_Δ1-100_* or pET28a::_7a1_*aopW1-7a1_Δ1-100_* (encoding His-tagged AopW1_101-485_ from strains M6 and 7a1, respectively) were grown in 5 ml of LB supplemented with kanamycin at 37 °C overnight. Cultures were then transferred to 100 ml of the same medium and incubated at 37 °C until the OD_600_ reached to 0.6. Expression was induced by addition of 1 mM isopropyl β-d-1-thiogalactopyranoside (IPTG). After 4 h, cells were resuspended in NPI buffer (50 mM NaH_2_PO_4_, 300 mM NaCl; pH 8.0), containing 10 mM imidazole (BioWorld, USA), harvested by centrifugation at 6,000 *g* for 15 min at 4 °C, and stored at −80 °C until use.

For purification of recombinant proteins, pellets from 100 ml of IPTG-induced cultures were resuspended in 3 ml of NPI buffer containing 1 mM imidazole, 1 mM phenylmethanesulfonyl fluoride (PMSF), 1X protease inhibitor cocktail (Bimake, USA) and 1 mg/ml lysozyme. Suspensions were incubated on ice for 30 min, and then sonicated on ice ten times for 15 s, with 15-s cooling intervals between sonication treatments. The lysate was clarified by centrifugation at 10,000 *g* (30 min, at 4 °C). Half of the clarified lysate was incubated at 4 °C for 90 min in a column containing 130 μl of Protino Ni-NTA agarose (Macherey Nagel), previously equilibrated with NPI buffer (pH 8.0), containing 10 mM imidazole, allowing binding of His-tagged AopW1 to Protino Ni-NTA agarose. Columns were washed twice with 500 μl of NPI buffer containing 100 mM imidazole, and once with 500 μl of NPI buffer containing 200 mM imidazole. His-tagged AopW1 was eluted by adding twice 50 μl of NPI buffer containing 500 mM imidazole. Purified proteins were dialyzed using 10,000 MWCO Slide-A-Lyzer Dialysis Cassettes (Thermo Fisher Scientific) and concentrated using Amicon Ultra-4 Centrifugal Filter Unit (Millipore Sigma, USA), following the manufacturer’s instructions. Expression and purification of His-tagged AopW1 were verified by sodium dodecyl sulfate-polyacrylamide gel electrophoresis (SDS-PAGE) and confirmed by Western blot using His-tag monoclonal antibodies.

### Non-muscle F-actin disruption assays

F-actin disruption assays were carried out using the Actin Binding Protein Spin-Down Assay Biochem kit (Cytoskeleton, USA) following manufacturer’s instructions, in the presence of purified recombinant AopW1_101-485_. Non-muscle actin was polymerized to actin filaments (F-actin) in 10 mM Tris pH 7.0, 1 mM ATP, 50 mM KCl, 1 mM EGTA, 0.2 mM CaCl_2_ and 2 mM MgCl_2_ for 1.5 h at 24 °C. Then, 10 mM of preassembled F-actin were incubated with 35 μg of AopW1_101-485_, 2 μM α-actinin (positive control) or 2 μM BSA (negative control) for 1 h at 24 °C and centrifuged at 150,000 *g* for 1.5 h. Proteins from 20-μl aliquots from pellet or supernatant fractions were separated in a 4-20% SDS-PAGE gel and stained with Coomassie blue solution (Fermentas). To check the presence of AopW1_101-485_ in the supernatant fraction, Western blots were performed using His-Tag monoclonal antibodies.

### *Agrobacterium-mediated* transient expression on *N. benthamiana* leaves and assessment of subcellular localization

Transient expression experiments were carried out as described by Roden *et al*. (2004) with some modifications. Briefly, overnight cultures of *A. tumefaciens* GV3101 carrying the different plasmids were washed with a 10 mM MgCl_2_ solution, centrifuged at 3,500 *g* for 5 min and resuspended in induction solution containing 10 mM MgCl_2_, 200 mM acetosyringone and 10 mM 2-(N-morpholino)-ethanesulfonic acid (MES) (pH 5.6). The suspensions were then incubated at 25 °C without shaking for 3 h. Bacterial cultures were diluted to OD_600_ of 0.6 and infiltrated with a needleless syringe into the abaxial part of leaves of 4-week-old *N. benthamiana* plants.

Subcellular localization of AopW1 was assessed 24 and 48 h after inoculation with *A. tumefaciens* GV3101 carrying plasmid pEarleyGate 101 with *aopW1* fused in frame with the yellow fluorescent protein (YFP) gene and a HA tag, and the same strain carrying vectors with different plant markers, respectively. The tested markers were: *Discosoma* red fluorescent protein (DsRed) fused in frame with the actin binding domain 2 (ABD2) (DsRed-ABD2; Voigt *et al*., 2005b); monomeric red fluorescence protein (mRFP) fused in frame with the ER marker HDEL (mRFP-HDEL; Runions *et al*., 2006; Schoberer *et al*., 2009); monomeric Cherry Fluorescent Protein (mCherry) fused in frame with the endosome marker Wave33 (RabD2b) (Bar *et al*., 2013); the endosome marker FYVE fused in frame with DsRed (Voigt *et al*., 2005a); and the endosome markers Ara7 (RabF2b) and AtEHD1 fused in frame with cyan fluorescent protein (CFP) (Bar *et al*., 2008a; Bar *et al*., 2013; Lee *et al*., 2004). Alternatively, samples were stained with 1 mg/ml 4’,6-diamidino-2-phenylindole (DAPI) for detection of the plant cell nucleus. Detection with infrared emission was used to locate the chloroplasts. *Agrobacterium tumefaciens* GV3101 derivative strains carrying pEarleyGate 104 and pEarleyGate 100 were used as positive and negative controls, respectively. Functional fluorophores were visualized using a SPE confocal microscope (Leica Microsystems). Contrast and intensity for each image were manipulated uniformly using ImageJ software (Schneider *et al*., 2012). Experiments were carried out at least twice for each effector/marker combination. Expression of AopW1-YFP on *N. benthamiana* leaves was validated by Western blot analysis using an HA primary antibody.

### Hypersensitive response (HR) and ethylene biosynthesis assays in *N. tabacum* leaves

For HR assays, leaves of 5-6-week-old *N. tabaccum* cv. Samsun NN were transiently transformed with *A. tumefaciens* GV3101 harboring *35S::TvEIX* and *35S::EHD1-GFP* or *35S::free-GFP* (mock). HR development was quantified 72 h post injection using ImageJ. Plants were kept in a greenhouse under long day conditions (16h light/ 8h dark), at 25 °C. Ethylene biosynthesis was measured as previously described by Leibman-Markus *et al*. (2017). Briefly, leaf disks (0.9 cm diameter) were taken from EHD1-GFP transiently expressing *N. tabaccum* cv. Samsun NN plants (same construct as described above), 40 h post injection. Free-GFP (empty vector) was injected as mock. Five disks were sealed in each 10 ml flasks containing 1 ml assay medium (with or without 1 μg/ml EIX) and incubated with shaking for 4 h at room temperature. Ethylene production was measured by Gas chromatography (Varian 3350, Varian, USA).

### Detection of callose deposition in *N. benthamiana* leaves

Callose deposition assays were carried out by the procedure described by Nguyen *et al*. (2010), with some modifications. Briefly, *N. benthamiana* leaves were infiltrated with 40 μM flg22 (Felix *et al*., 1999) with a needleless syringe. After 24 h, treated leaves were agroinfiltrated with *A. tumefaciens* GV3101 strain carrying pEarleyGate 101 vector, either empty or containing *aopW1* fused to the YFP gene (OD600 adjusted to 0.6). After 24 h, 1-cm diameter disks were collected and serially incubated in 95%, 70% and 50% ethanol solutions, for 6, 12 and 2 h, respectively, at 37 °C with shaking. The ethanol solutions were replaced several times. Then, disks were washed with double distilled water (DDW) and stained for 2 h with 1% aniline blue in 150 mM K2HPO4 (pH 9.5). Disks were then transferred to a slide containing 60% glycerol in PBS, and callose deposits were visualized using a SPE confocal microscope using 405 nm laser for aniline blue excitation and a 475-525 nm band-pass emission filter for aniline blue fluorescence collection. Contrast and intensity for each image were manipulated uniformly using ImageJ and deposits were counted from 6 areas from 3 different leaves from 3 plants (54 total areas per experiment).

## SUPPLEMENTARY MATERIAL

Supplementary material is provided in a separate pdf file.

## ACKNOWLEDGEMENTS

This work was supported by research grant IS-5023-17C from the United States-Israel Binational Agriculture Research and Development (BARD) Fund. FPM was recipient of the José Castillejo grant of the Ministry of Education, Culture and Sport of the Spanish Government and VI PPIT-US grant of the University of Seville. GMS was recipient of a Lady Davis post-doctoral fellowship. We thank Dr. Joanna Jelenska and Prof. Jean T. Greenberg for kindly providing the pBAV154-dexHopW1-HA plasmid that served as template for amplification of the *P. syringae hopW1* ORF, and Dr. Johannes Sikorski for providing genomic DNA of *X. translucens* pv. *translucens* DSM 18974 that was used for amplification of the ORF of the *hopW1* homolog of this strain. We also thank Prof. Nahum Shpigel for his advices regarding actin staining in yeast, and Dr. Einat Zelinger for the help provided in microscopy techniques.

