## Supplementary material for "Natural variation in a short region of the *Acidovorax citrulli* type III-secreted effector AopW1 is associated with differences in cytotoxicity"

**Contents** **Page**

Supplementary Figure S1……………………………………………………….............. 2

Supplementary Figure S2…………………………………………………..………... 3

Supplementary Figure S3…………………………………………………..………... 4

Supplementary Table S1……………………………………………………….................5

Supplementary Table S2……………………………………………………….................6

Supplementary Table S3……………………………………………………….................13

Supplementary Table S4……………………………………………………….................15

Supplementary Appendix S1…………………………………………………….......... 20

References of Supplementary Material……………………………………………. 25

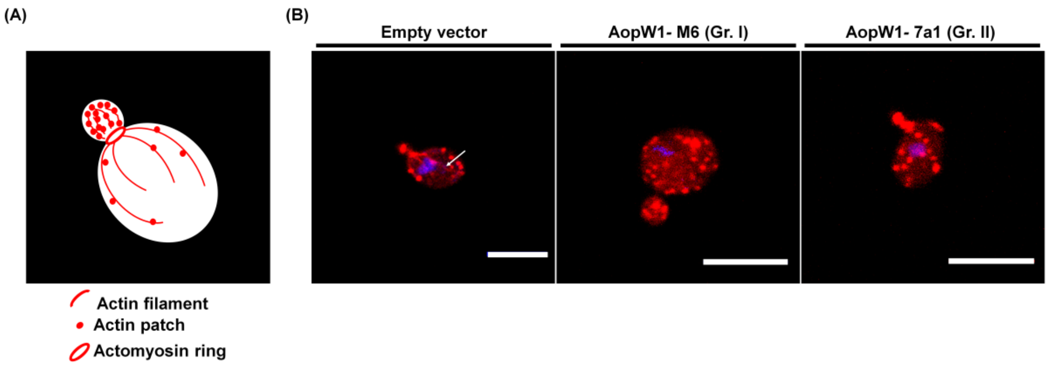


**Supplementary Figure S1**. **Group I AopW1 disrupts actin filaments in yeast. (A)** Scheme of the regular actin organization during mitosis in budding yeast. F-actin forms three distinct filamentous structures: actin filaments, actin patches and the actin ring. Actin filaments run along the cell and actin patches distribution is correlated with the polarized growth region. **(B)** Actin staining in yeast expressing *aopW1* from strains M6 (group I) or 7a1 (group II) or carrying empty vector. The yeast nucleus is coloured with blue (DAPI). Actin cables and patches are coloured with red (phalloidin-TRITC). White arrow points an actin cable (F-actin). Scale bar = 5 µm. The experiment was performed three times with similar results.

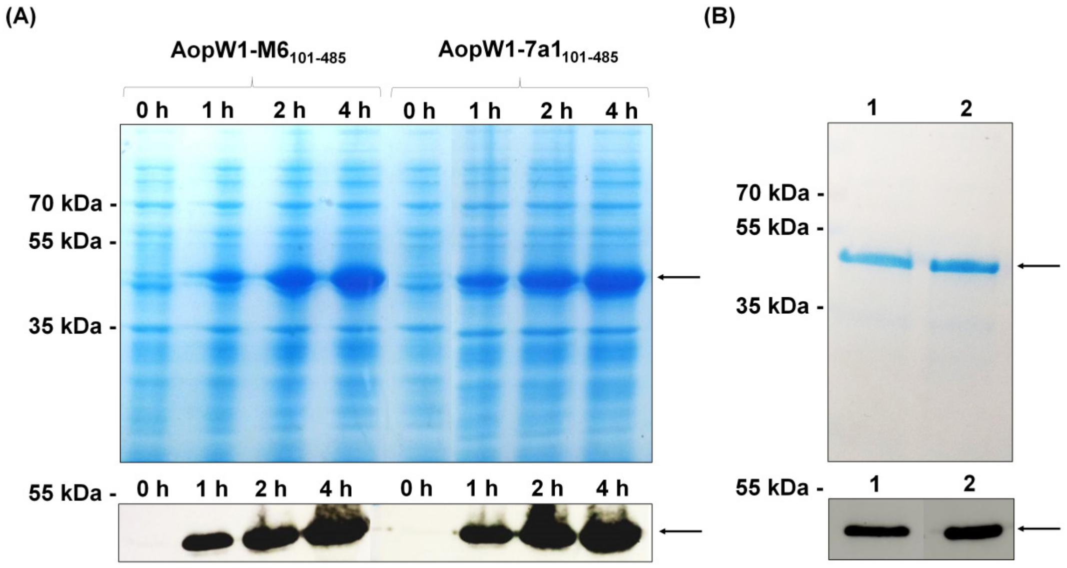


**Supplementary Figure S2**. ***In vitro* expression and purification of recombinant AopW1$_{101\text{-}485}$ proteins in *Escherichia coli*.** (**A**) Expression of AopW1$_{101\text{-}485}$ from M6 (group I) and 7a1 (group II) *A. citrulli* strains in *E. coli* BL21(DE3) at 0, 1, 2 and 4 h after induction by 1 mM IPTG. **(B)** Purification of AopW1$_{101\text{-}485}$ from strains M6 and 7a1 4 h after induction by 1 mM IPTG. Lines 1 and 2 show purified AopW1$_{101\text{-}485}$ from M6 and 7a1, respectively. Upper panels: Coomassie blue staining. Bottom panels: immunodetection of AopW1$_{101\text{-}485}$ using an anti-His primary antibody. Black arrows indicate AopW1$_{101\text{-}485}$.

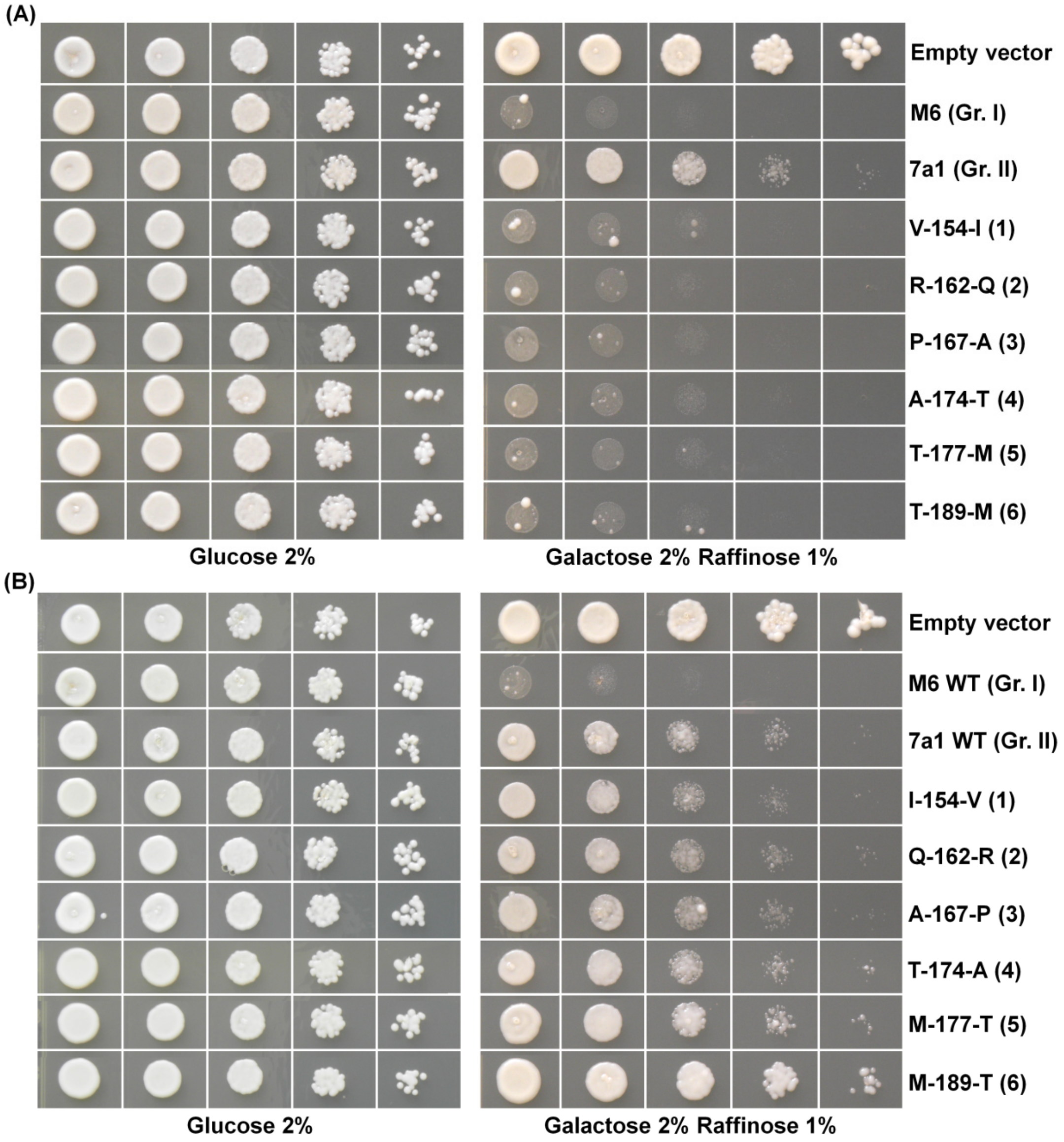


**Supplementary Figure S3. Effects of single amino acid substitutions in the AopW1 HVR on yeast growth inhibition. (A)** AopW1 variants of strain M6 (group I) carrying single substitutions. **(B)** AopW1 variants of strain 7a1 (group II) carrying single substitutions. *Saccharomyces cerevisiae* BY4741 carrying the different *aopW1* variants cloned into plasmid pGML10, or the empty vector, were grown in glucose (repressing) and in galactose + raffinose (inducing) medium. Pictures were taken after 3 days of growth at 28 °C and are representative of three independent experiments. See Supplementary Table S2 for variant details.

**Supplementary Table S1.** Yeast growth-inhibition phenotypes[1] induced by different versions of *Acidovorax citrulli* AopW1 and homolog effectors under regular conditions or under stress.

| **Effector** | **Strain/group** | **No stressors** | **Caffeine** | **NaCl** | **Sorbitol** |
|---|---|---|---|---|---|
| AopW1-M6 | *A. citrulli* M6 (Gr. I) | + + + | + + + | + + + + | + + + |
| AopW1-7a1 | *A. citrulli* 7a1 (Gr. II) | + | + + | + + | + + |
| HopW1 | *P. syringae* pv. *maculicola* ES4326 | + + + | + + + + | + + + + | + + + + |
| HopW1 homolog | *X. translucens* DSM 18974 | + + | + + | + + + + | + + |
| Empty vector | | - | - | - | - |

[1]Degree of growth inhibition: -, no inhibition; + slight inhibition (growth up to $10^3$ dilution); + +, moderate inhibition (growth to up to $10^2$ dilution); + + +, strong inhibition (growth to up to $10^1$ dilution); + + + +, no growth. *S. cerevisiae* BY4741 with plasmid pGML10, either empty or carrying the T3E genes, were grown in glucose (repressing) and in galactose + raffinose (inducing) medium. Growth-inhibition phenotypes were observed after 3 days of growth at 28 °C. Results were displayed from phenotypes observed in inducing medium compared to those from repressing medium.

**Supplementary Table S2.** Strains and plasmids used in this study.

| Strain/plasmid | Relevant properties | Source or reference |
|---|---|---|
| ***Acidovorax citrulli*** | | |
| M6 | Wild-type, group I strain | Burdman *et al.*, 2005 |
| 7a1 | Wild-type, group II strain | Eckshtain-Levi *et al.*, 2014 |
| ***Escherichia coli*** | | |
| DB3.1 | *gyrA462*, *endA1*, Δ(*sr1-recA*), *mcrB*, *mrr*, *hsdS20*, *glnV44* (=*supE44*), *ara14*, *galK2*, *lacY1*, *proA2*, *rpsL20*, *xyl5*, *leuB6*, *mtl1* | Invitrogen, USA) |
| DH5α | *supE44*, Δ*lacU169*, *hsdR17*, *recA1*, *endA1*, *gyrA96*, *thi-1*, *relA1*, $Nx^R$ | Sambrook *et al.*, 1989 |
| BL21(DE3) | *E. coli* B $F^-$*dcmompThsdS* ($r_B^- m_B^-$) *gallon*λ (DE3 [*lacIlac*UV5-T7 gene *1ind1sam7nin5*]) | Studier and Moffatt, 1986 |
| S17-1 λpir | Δlysogenic S17-1 derivate producing π protein for replication of plasmids carrying *oriR6K*; *recA*, *pro*, *hsdR,* RP4-2-Tc::Mu-Km::Tn7, λ-pir | Simon *et al.*, 1983 |
| ***Agrobacterium tumefaciens*** | | |
| GV3101 | wild type; $Rif^R$ | Rotino and Gleddie, 1990 |
| ***Saccharomyces cerevisiae*** | | |
| BY4741 | *MATa, his3Δ1, leu2Δ0, met15Δ0, ura3Δ0* | Brachmann *et al.*, 1998 |
| **Plasmids** | | |
| pGML10 | *E. coli-S. cerevisiae* shuttle vector, 9E10 epitope (Myc-tag), centromere-type ori, GAL1-10 promoter, $Ap^R$, LEU2 | Iha and Tsurugi, 1998 |
| pET28a(+) | Plasmid with His N and C terminal tag for bacterial expression; $Km^R$ | Novagen (Germany) |
| pDONR207 | Gateway donor vector; $Gm^R$ | Invitrogen |
| pEarleyGate 100 | Gateway-compatible plant transformation vector; $Km^R$ | Earley *et al.*, 2006 |
| pEarleyGate 101 | Gateway-compatible plant transformation vector with YFP and HA C-terminal tags; $Km^R$ | Earley *et al.,* 2006 |
| pEarleyGate 104 | Gateway-compatible plant transformation vector with a YFP N-terminal tag; $Km^R$ | Earley *et al.*, 2006 |

| | | |
|---|---|---|
| pBAV154-dexHopW1-HA | Gateway binary plant expression vector carrying the open reading frame (ORF) of *hopW1* from *Pseudomonas syringae* pv. *maculicola* strain ES4326; used as template for cloning into pGML10; $Km^R$; $Cm^R$ | Kang *et al.*, 2014 |
| pGML10::$_{Ps}$*hopW1* | pGML10 containing the ORF of *hopW1* from *Pseudomonas syringae* pv. *maculicola* strain ES4326; $Ap^R$; LEU2 | This study |
| pGML10::$_{Xt}$*hopW1* | pGML10 containing the ORF of the *hopW1* homolog from *Xanthomonas translucens* pv. *translucens* strain DSM 18974; $Ap^R$; LEU2 | This study |
| pGML10::$_{M6}$*aopW1* | pGML10 containing the complete ORF of the *aopW1* (*APS58_3289*) gene of *A. citrulli* M6 (group I); $Ap^R$; LEU2 | This study |
| pGML10::$_{7a1}$*aopW1* | pGML10 containing the complete ORF of the *aopW1* gene of *A.citrulli* 7a1 (group II); $Ap^R$; LEU2 | This study |
| pGML10::$_{M6}$*aopW1*$_{V154I}$ | pGML10 carrying the M6 *aopW1* ORF in which the amino acid (a.a.) in position 154 (V; position 1) was replaced by the corresponding a.a. present in the 7a1 strain (I); $Ap^R$; LEU2 | This study |
| pGML10::$_{M6}$*aopW1*$_{R162Q}$ | pGML10 carrying the M6 *aopW1* ORF in which the a.a. in position 162 (R; position 2) was replaced by the corresponding a.a. present in the 7a1 strain (Q); $Ap^R$; LEU2 | This study |
| pGML10::$_{M6}$*aopW1*$_{P167A}$ | pGML10 carrying the M6 *aopW1* ORF in which the a.a. in position 167 (P; position 3) was replaced by the corresponding a.a. present in the 7a1 strain (A); $Ap^R$; LEU2 | This study |
| pGML10::$_{M6}$*aopW1*$_{A174T}$ | pGML10 carrying the M6 *aopW1* ORF in which the a.a. in position 174 (A; position 4) was replaced by the corresponding a.a. present in the 7a1 strain (T); $Ap^R$; LEU2 | This study |
| pGML10::$_{M6}$*aopW1*$_{T177M}$ | pGML10 carrying the M6 *aopW1* ORF in which the a.a. in position 177 (T; position 5) was replaced by the corresponding a.a. present in the 7a1 strain (M); $Ap^R$; LEU2 | This study |

| | | |
|---|---|---|
| pGML10::M6*aopW1*T189M | pGML10 carrying the M6 *aopW1* ORF in which the a.a. in position 189 (T; position 6) was replaced by the corresponding a.a. present in the 7a1 strain (M); $Ap^R$; LEU2 | This study |
| pGML10::M6*aopW1*L319S | pGML10 carrying the M6 *aopW1* ORF in which the a.a. in position 319 (L) was replaced by the corresponding a.a. present in the 7a1 strain (S); $Ap^R$; LEU2 | This study |
| pGML10::M6*aopW1*1+2 | pGML10 carrying the M6 *aopW1* ORF in which the a.a. in positions 154 and 162 were replaced by the corresponding a.a. present in the 7a1 strain; $Ap^R$; LEU2 | This study |
| pGML10::M6*aopW1*1+2+3 | pGML10 carrying the M6 *aopW1* ORF in which the a.a. in positions 154, 162 and 167 were replaced by the corresponding a.a. present in the 7a1 strain; $Ap^R$; LEU2 | This study |
| pGML10::M6*aopW1*1+2+3+4 | pGML10 carrying the M6 *aopW1* ORF in which the a.a. in positions 154, 162, 167 and 174 were replaced by the corresponding a.a. present in the 7a1 strain; $Ap^R$; LEU2 | This study |
| pGML10::M6*aopW1*1+2+3+4+6 | pGML10 carrying the M6 *aopW1* ORF in which the a.a. in positions 154, 162, 167, 174 and 189 were replaced by the corresponding a.a. present in the 7a1 strain; $Ap^R$; LEU2 | This study |
| pGML10::M6*aopW1*1+2+3+6 | pGML10 carrying the M6 *aopW1* ORF in which the a.a. in positions 154, 162, 167 and 189 were replaced by the corresponding a.a. present in the 7a1 strain; $Ap^R$; LEU2 | This study |
| pGML10::M6*aopW1*1+6 | pGML10 carrying the M6 *aopW1* ORF in which the a.a. in positions 154 and 189 were replaced by the corresponding a.a. present in the 7a1 strain; $Ap^R$; LEU2 | This study |
| pGML10::7a1*aopW1*I154V | pGML10 carrying the 7a1 *aopW1* ORF in which the a.a. in position 154 (I; position 1) was replaced by the corresponding a.a. present in the M6 strain (V); $Ap^R$; LEU2 | This study |
| pGML10::7a1*aopW1*Q162R | pGML10 carrying the 7a1 *aopW1* ORF in which the a.a. in position 162 (Q; position 2) was replaced by the corresponding a.a. present in the M6 strain (R); $Ap^R$; LEU2 | This study |

| | | |
|---|---|---|
| pGML10::$_{7a1}$*aopW1*$_{A167P}$ | pGML10 carrying the 7a1 *aopW1* ORF in which the a.a. in position 167 (A; position 3) was replaced by the corresponding a.a. present in the M6 strain (P); Ap$^{R}$; LEU2 | This study |
| pGML10::$_{7a1}$*aopW1*$_{T174A}$ | pGML10 carrying the 7a1 *aopW1* ORF in which the a.a. in position 174 (T; position 4) was replaced by the corresponding a.a. present in the M6 strain (A); Ap$^{R}$; LEU2 | This study |
| pGML10::$_{7a1}$*aopW1*$_{M177T}$ | pGML10 carrying the 7a1 *aopW1* ORF in which the a.a. in position 177 (M; position 5) was replaced by the corresponding a.a. present in the M6 strain (T); Ap$^{R}$; LEU2 | This study |
| pGML10::$_{7a1}$*aopW1*$_{M189T}$ | pGML10 carrying the 7a1 *aopW1* ORF in which the a.a. in position 189 (M; position 6) was replaced by the corresponding a.a. present in the M6 strain (T); Ap$^{R}$; LEU2 | This study |
| pGML10::$_{7a1}$*aopW1*$_{S319L}$ | pGML10 carrying the 7a1 *aopW1* ORF in which the a.a. in position 319 (S) was replaced by the corresponding a.a. present in the M6 strain (L); Ap$^{R}$; LEU2 | This study |
| pGML10::$_{7a1}$*aopW1*$_{1+2}$ | pGML10 carrying the 7a1 *aopW1* ORF in which the a.a. in positions 154 and 162 were replaced by the corresponding a.a. present in the M6 strain; Ap$^{R}$; LEU2 | This study |
| pGML10::$_{7a1}$*aopW1*$_{1+2+3}$ | pGML10 carrying the 7a1 *aopW1* ORF in which the a.a. in positions 154, 162 and 167 were replaced by the corresponding a.a. present in the M6 strain; Ap$^{R}$; LEU2 | This study |
| pGML10::$_{7a1}$*aopW1*$_{1+2+3+4}$ | pGML10 carrying the 7a1 *aopW1* ORF in which the a.a. in positions 154, 162, 167 and 174 were replaced by the corresponding a.a. present in the M6 strain; Ap$^{R}$; LEU2 | This study |
| pGML10::$_{7a1}$*aopW1*$_{1+2+3+4+6}$ | pGML10 carrying the 7a1 *aopW1* ORF in which the a.a. in positions 154, 162, 167, 174 and 189 were replaced by the corresponding a.a. present in the M6 strain; Ap$^{R}$; LEU2 | This study |
| pGML10::$_{7a1}$*aopW1*$_{1+2+6}$ | pGML10 carrying the 7a1 *aopW1* ORF in which the a.a. in positions 154, 162 and 189 were replaced by the corresponding a.a. present in the M6 strain; Ap$^{R}$; LEU2 | This study |

| | | |
|---|---|---|
| pGML10::7a1*aopW1*1+6 | pGML10 carrying the 7a1 *aopW1* ORF in which the a.a. in positions 154 and 189 were replaced by the corresponding a.a. present in the M6 strain; $Ap^R$; LEU2 | This study |
| pGML10::M6*aopW1*Δ1-50 | pGML10 carrying the M6 *aopW1* ORF without the first 150 bp; $Ap^R$; LEU2 | This study |
| pGML10:: M6*aopW1*Δ1-135 | pGML10 carrying the M6 *aopW1* ORF without the first 405 bp; $Ap^R$; LEU2 | This study |
| pGML10::M6*aopW1*Δ1-145 | pGML10 carrying the M6 *aopW1* ORF without the first 435 bp; $Ap^R$; LEU2 | This study |
| pGML10:: M6*aopW1*Δ1-175 | pGML10 carrying the M6 *aopW1* ORF without the first 525 bp; $Ap^R$; LEU2 | This study |
| pGML10::M6*aopW1*Δ476-485 | pGML10 carrying the M6 *aopW1* ORF without the last 30 bp; $Ap^R$; LEU2 | This study |
| pGML10::M6*aopW1*Δ461-485 | pGML10 carrying the M6 *aopW1* ORF without the last 75 bp; $Ap^R$; LEU2 | This study |
| pGML10::M6*aopW1*Δ436-485 | pGML10 carrying the M6 *aopW1* ORF without the last 150 bp; $Ap^R$; LEU2 | This study |
| pET28a::M6*aopW1*Δ1-100 | pET28a carrying the M6 *aopW1* ORF lacking the first 300 bp, tagged with His in both extremes; used for *in vitro* actin depolymerization assays; $Km^R$ | This study |
| pET28a::7a1*aopW1*Δ1-100 | pET28a carrying the 7a1 *aopW1* ORF lacking the first 300 bp sequence, tagged with His in both extremes, used for *in vitro* actin depolymerization assays; $Km^R$ | This study |
| pDONR207::M6*aopW1* | pDONR207 carrying the M6 *aopW1* ORF without the stop codon (1455 bp); $Gm^R$ | This study |
| pEarleyGate 101::M6*aopW1* | pEarleyGate 101 carrying the M6 *aopW1* ORF fused to YFP and HA; $Km^R$ | This study |
| pDONR207::M6*aopW1*1+2+3 | pDONR207 carrying the M6 *aopW1* ORF in which the a.a. in positions 154, 162 and 167 were replaced by the corresponding a.a. present in the 7a1 strain without the stop codon (1455 bp); $Gm^R$ | This study |
| pEarleyGate 101::M6*aopW1*1+2+3 | pEarleyGate 101 carrying the M6 *aopW1* ORF in which the a.a. in positions 154, 162 and 167 were replaced by the corresponding a.a. present in the 7a1, fused to YFP and HA; $Km^R$ | This study |
| pDONR207:: 7a1*aopW1* | pDONR207 carrying the 7a1 *aopW1* ORF without the stop codon (1455 bp); $Gm^R$ | This study |

| | | |
|---|---|---|
| pEarleyGate 101::$_{7a1}$*aopW1* | pEarleyGate 101 carrying the 7a1 *aopW1* ORF fused to YFP and HA; $Km^R$ | This study |
| pDONR207:: $_{7a1}$*aopW1*$_{1+2+3+4}$ | pDONR207 carrying the 7a1 *aopW1* ORF in which the a.a. in positions 154, 162, 167 and 174 were replaced by the corresponding a.a. present in strain M6 without the stop codon (1455 bp); $Gm^R$ | This study |
| pEarleyGate 101::$_{7a1}$*aopW1*$_{1+2+3+4}$ | pEarleyGate 101 carrying the 7a1 *aopW1* ORF in which the a.a. in positions 154, 162, 167 and 174 were replaced by the corresponding a.a. present in strain M6, fused to YFP and HA; $Km^R$ | This study |
| pDONR207:: $_{7a1}$*aopW1*$_{\Delta 1\text{-}85}$ | pDONR207 carrying the 7a1 *aopW1* lacking the first 255 bp without the stop codon (1200 bp); $Gm^R$ | This study |
| pEarleyGate 101::$_{7a1}$*aopW1*$_{\Delta 1\text{-}85}$ | pEarleyGate 101 carrying the 7a1 *aopW1* ORF lacking the first 255 bp, fused to YFP and HA; $Km^R$ | This study |
| DsRed-ABD2 vector | Plasmid used for transient expression of the actin marker Actin Binding Domain 2 (ABD2) fused to the DsRed fluorescent protein; $Km^R$ | Voigt *et al.*, 2005b |
| mRFP-HDEL vector | Plasmid used for transient expression of the endoplasmic reticulum marker HDEL fused to red fluorescent protein; $Km^R$ | Runions *et al.*, 2006; Schoberer *et al.*, 2009 |
| DsRed-FYVE vector | Plasmid used for transient expression of the plant endosome marker FYVE fused to the DsRed fluorescent protein; $Km^R$ | Voigt *et al.*, 2005a |
| mCherry-Wave33 vector | Plasmid used for transient expression of the plant endosome marker Wave33 (RabD2b) fused to the mCherry fluorescent protein; $Km^R$ | Bar *et al.*, 2013 |
| mCherry-Wave34 vector | Plasmid used for transient expression of the plant endosome marker Wave34 (RabA1e) fused to the mCherry fluorescent protein; $Km^R$ | Bar *et al.*, 2013 |
| AtEHD1-CFP vector | Plasmid used for transient expression of the plant endosome marker AtEHD1 fused to the CFP fluorescent protein; $Km^R$ | Bar *et al.*, 2008 |
| Ara7-CFP vector | Plasmid used for transient expression of the plant endosome marker Ara7 fused to the CFP fluorescent protein; $Km^R$ | Lee *et al.*, 2004; Bar *et al.*, 2013 |

| | | |
|---|---|---|
| *Pro35S*:AtEHD1-GFP | Plasmid used for transient expression of the plant endosome marker AtEHD1 fused to the GFP fluorescent protein; $Km^R$ | Bar *et al*., 2008 |
| *Pro35S*:TvEIX | Plasmid used for transient expression of the *Trichoderma* derived ethylene inducing xylanase (EIX); $Km^R$ | Ron and Avni, 2004 |

* $Ap^R$, $Cm^R$, $Gm^R$, $Km^R$ and $Rif^R$ indicate resistance to ampicillin, chloramphenicol, gentamicin, kanamycin and rifampicin, respectively.

**Supplementary Table S3.** Details of the amino acid residues and positions subjected to site-directed mutagenesis in the HVR of *A. citrulli* M6 (group I) and 7a1 (group II) AopW1.

| M6 AopW1 HVR variants[1] | Amino acid position (base pair in ORF)[2] | Native codon in M6 | Changed codon in M6 |
|---|---|---|---|
| 1 | V-154-I (460) | GTC (V) | ATC (I) |
| 2 | R-162-Q (485-486) | CGT (R) | CAA (Q) |
| 3 | P-167-A (499) | CCC (P) | GCC (A) |
| 4 | A-174-T (520) | GCC (A) | ACC (T) |
| 5 | T-177-M (530) | ACG (T) | ATG (M) |
| 6 | T-189-M (566) | ACG (T) | ATG (M) |
| 1+2 | V-154-I (460)<br>R-162-Q (485-486) | GTC (V)<br>CGT (R) | ATC (I)<br>CAA (Q) |
| 1+2+3 | V-154-I (460)<br>R-162-Q (485-486)<br>P-167-A (499) | GTC (V)<br>CGT (R)<br>CCC (P) | ATC (I)<br>CAA (Q)<br>GCC (A) |
| 1+2+3+4 | V-154-I (460)<br>R-162-Q (485-486)<br>P-167-A (499)<br>A-174-T (520) | GTC (V)<br>CGT (R)<br>CCC (P)<br>GCC (A) | ATC (I)<br>CAA (Q)<br>GCC (A)<br>ACC (T) |
| 1+2+3+4+6 | V-154-I (460)<br>R-162-Q (485-486)<br>P-167-A (499)<br>A-174-T (520)<br>T-189-M (566) | GTC (V)<br>CGT (R)<br>CCC (P)<br>GCC (A)<br>ACG (T) | ATC (I)<br>CAA (Q)<br>GCC (A)<br>ACC (T)<br>ATG (M) |
| 1+2+3+6 | V-154-I (460)<br>R-162-Q (485-486)<br>P-167-A (499)<br>T-189-M (566) | GTC (V)<br>CGT (R)<br>CCC (P)<br>ACG (T) | ATC (I)<br>CAA (Q)<br>GCC (A)<br>ATG (M) |
| 1+6 | V-154-I (460)<br>T-189-M (566) | GTC (V)<br>ACG (T) | ATC (I)<br>ATG (M) |
| **7a1 AopW1 HVR variants[1]** | **Amino acid position (base pair in ORF)[2]** | **Native codon in 7a1** | **Changed codon in 7a1** |
| 1 | I-154-V (460) | ATC (I) | GTC (V) |
| 2 | Q-162-R (485-486) | CAA (Q) | CGT (R) |
| 3 | A-167-P (499) | GCC (A) | CCC (P) |
| 4 | T-174-A (520) | ACC (T) | GCC (A) |
| 5 | M-177-T (530) | ATG (M) | ACG (T) |
| 6 | M-189-T (566) | ATG (M) | ACG (T) |

| | | | |
|---|---|---|---|
| 1+2 | I-154-V (460)<br>Q-162-R (485-486) | ATC (I)<br>CAA (Q) | GTC (V)<br>CGT (R) |
| 1+2+3 | I-154-V (460)<br>Q-162-R (485-486)<br>A-167-P (499) | ATC (I)<br>CAA (Q)<br>GCC (A) | GTC (V)<br>CGT (R)<br>CCC (P) |
| 1+2+3+4 | I-154-V (460)<br>Q-162-R (485-486)<br>A-167-P (499)<br>T-174-A (520) | ATC (I)<br>CAA (Q)<br>GCC (A)<br>ACC (T) | GTC (V)<br>CGT (R)<br>CCC (P)<br>GCC (A) |
| 1+2+3+4+6 | I-154-V (460)<br>Q-162-R (485-486)<br>A-167-P (499)<br>T-174-A (520)<br>M-189-T (566) | ATC (I)<br>CAA (Q)<br>GCC (A)<br>ACC (T)<br>ATG (M) | GTC (V)<br>CGT (R)<br>CCC (P)<br>GCC (A)<br>ACG (T) |
| 1+2+6 | I-154-V (460)<br>Q-162-R (485-486)<br>M-189-T (566) | ATC (I)<br>CAA (Q)<br>ATG (M) | GTC (V)<br>CGT (R)<br>ACG (T) |
| 1+6 | I-154-V (460)<br>M-189-T (566) | ATC (I)<br>ATG (M) | GTC (V)<br>ACG (T) |

[1]M6/7a1 variants carry individual or combined substitutions in positions 1-6 as indicated in Figure 4B.

[2]The position(s) of the substituted nucleotide(s) in the ORF is/are indicated between parentheses.

**Supplementary Table S4.** DNA oligonucleotide primers used in this study.

| **Name** | **Sequence**[2] | **Use** |
|---|---|---|
| pGML10_F | 5'-ACTTTCAACATTTTCGGTTTGT-3' | Sequence verification of inserts cloned into pGML10 |
| pGML10_R | 5'-ATTCGCCCGGAATTAGCTTG-3' | |
| pDONR_F | 5'-CGTTAACGCTAGCATGGATCTC-3' | Sequence verification of inserts cloned into pDONR207 |
| pDONR_R | 5'-GTAACATCAGAGATTTTGAGAC-3' | |
| Psm_HopW1__F[1] | 5'-GGAGAAAAAACCCCGGATCGAATTGACTATGAGTCCAGCTCAGATTATCCGCAC-3' | Used to amplify the open reading frame (ORF) of *hopW1* from *Pseudomonas syringae* pv. *maculicola* ES4326 for expression in yeast |
| Psm_HopW1_R[1] | 5'-AGATCCTCTTCGGAAATCAGTTTCTGTTCAGAACGGTCTTTGGAGGACTTGTTTGA-3' | |
| Xt_HopW1__F[1] | 5'-GGAGAAAAAACCCCGGATCGAATTGACTATGGGAGGTAGAATCACAAGAGA-3' | Used to amplify the ORF of the *hopW1* homolog gene of *Xanthomonas translucens* DSM 18974 for expression in yeast |
| Xt_HopW1_R[1] | 5'-AGATCCTCTTCGGAAATCAGTTTCTGTTCATAGGCTTTCTTTGAAGATGAATCG-3' | |
| AopW1_BamHI_F | 5'-GGGTCTAGAATGCCTCTACAGTCCATTTCC-3' | Used to amplify the complete ORF of M6 and 7a1 *aopW1* for expression in yeast |
| AopW1_EcoRI_R | 5'-CAGAATTCGATGGTTGATCCCCCGTC-3' | |
| M6AopW1V154I_F | 5'-CGGGCACGGCTCC**A**TCATTGATGTGCG-3 | Used to generate a mutated variant of M6 AopW1, switching the amino acid in position 154 from valine to isoleucine |
| M6AopW1V154I_RR | 5'-CGCACATCAATGA**T**GGAGCCGTGCCCG-3 | |
| M6AopW1R162Q_F | 5'-CGACGCCCTCCGC**AA**GGCTCCCGGATTC-3 | Used to generate a mutated variant of M6 AopW1, switching the amino acid in position 162 |

| | | |
|---|---|---|
| M6AopW1R162Q_RR | 5'-GAATCCGGGAGCC**TT**GCGGAGGGCGTCG-3 | from arginine to glutamine |
| M6AopW1P167A_F | 5'-CGTGGCTCCCGGATT**G**CCGAAGGC-3 | Used to generate a mutated variant of M6 AopW1, switching the amino acid in position 167 from proline to alanine |
| M6AopW1P167A_RR | 5'-GCCTTCGGCAATCCG**G**GAGCCACG-3 | |
| M6AopW1A174T_F | 5'-CAGGAACGGGAG**A**CCTTCGCCACGG-3 | Used to generate a mutated variant of M6 AopW1, switching the amino acid in position 174 from alanine to threonine |
| M6AopW1A174T_RR | 5'-CCGTGGCGAAGG**T**CTCCCGTTCCTG-3 | |
| M6AopW1T177M_F | 5'-GGAGGCCTTCGCCA**T**GGTGCTCGAAGA-3 | Used to generate a mutated variant of M6 AopW1, switching the amino acid in position 177 from threonine to methionine |
| M6AopW1T177M_RR | 5'-TCTTCGAGCACCAT**G**GCGAAGGCCTCC-3 | |
| M6AopW1T189M_F | 5'-CGCAACCGTGGTAAGGACA**T**GCTGCGAACG-3 | Used to generate a mutated variant of M6 AopW1, switching the amino acid in position 189 from threonine to methionine |
| M6AopW1T189M_RR | 5'-CGTTCGCAGCATGTCCTTA**C**CACGGTTGCG-3 | |
| M6AopW1L319S_F | 5'-ATGAACACCATGGAGT**C**GTGGCCGAAAACGACG-3 | Used to generate a mutated variant of M6 AopW1, switching the amino acid in position 319 from leucine to serine |
| M6AopW1L319S_RR | 5'-CGTCGTTTTCGGCCAC**G**ACTCCATGGTGTTCAT-3 | |
| 7a1AopW1I154V_F | 5'-CACGAATGGCGCC**G**TCATCGACGTGCG-3 | Used to generate a mutated variant of 7a1 AopW1, switching the amino acid in position 154 |

| | | |
|---|---|---|
| 7a1AopW1I154V_RR | 5'-CGCACGTCGATGA**C**GGCGCCATTCGTG-3 | from isoleucine to valine |
| 7a1AopW1Q162R_F | 5'-CGGCGACCTGAGC**GT**CCGTCCCGCATTG-3 | Used to generate a mutated variant of 7a1 AopW1, switching the amino acid in position 162 from glutamine to arginine |
| 7a1AopW1Q162R_RR | 5'-CAATGCGGGACGG**AC**GCTCAGGTCGCCG-3 | |
| 7a1AopW1A167P_F | 5'-CAACCGTCCCGCATT**C**CCGAGGGC-3 | Used to generate a mutated variant of 7a1 AopW1, switching the amino acid in position 167 from alanine to proline |
| 7a1AopW1A167P_RR | 5'-GCCCTCGGGAATGCG**G**GACGGTTG-3 | |
| 7a1AopW1T174A_F | 5'-CAGGAGCGGGAG**G**CCTTCGCCATGG-3 | Used to generate a mutated variant of 7a1 AopW1, switching the amino acid in position 174 from threonine to alanine |
| 7a1AopW1T174A_RR | 5'-CCATGGCGAAGG**C**CTCCCGCTCCTG-3 | |
| 7a1AopW1M177T_F | 5'-GGGAGACCTTCGCCA**C**GGTGCTCGAAGAG-3 | Used to generate a mutated variant of 7a1 AopW1, switching the amino acid in position 177 from methionine to threonine |
| 7a1AopW1M177T_RR | 5'-CTCTTCGAGCACCGT**G**GCGAAGGTCTCCC-3 | |
| 7a1AopW1M189T_F | 5'-GCCGGGGGAAGGACA**C**GCTGCGCC-3 | Used to generate a mutated variant of 7a1 AopW1, switching the amino acid in position 189 from methionine to threonine |
| 7a1AopW1M189T_RR | 5'-GGCGCAGCGTGTCCT**T**CCCCCGGC-3 | |
| 7a1AopW1S319L_F | 5'-ATGAACACCATGGAGT**T**GTGGCCGAAAACGACG-3 | Used to generate a mutated variant of 7a1 AopW1, |

| | | |
|---|---|---|
| 7a1AopW1S319L_RR | 5'-CGTCGTTTTCGGCCAC**A**ACTCCATGGTGTTCAT-3 | switching the amino acid in position 319 from serine to leucine |
| dNt50M6AopW1_XbaI_F | 5'-AAATCTAGAATGGGCGCAAGACAACTGCCC-3 | Used to generate a mutated variant of M6 AopW1, deleting the first 50 amino acids |
| M6AopW1_EcoRI_R | 5'-AAAGAATTCTGGTTGATCCCCCGTCCG-3 | |
| dNt135M6AopW1_XbaI_F | 5'-AAATCTAGAATGGTGAAAGACCAGGTCGCCACG-3 | Used to generate a mutated variant of M6 AopW1, deleting the first 135 amino acids |
| M6AopW1_EcoRI_R | 5'-AAAGAATTCTGGTTGATCCCCCGTCCG-3 | |
| dNt145M6AopW1_XbaI_F | 5'-AAATCTAGAATGAGCATGGACGGGCACGGC-3 | Used to generate a mutated variant of M6 AopW1, deleting the first 145 amino acids |
| M6AopW1_EcoRI_R | 5'-AAAGAATTCTGGTTGATCCCCCGTCCG-3 | |
| dNt175M6AopW1_XbaI_F | 5'-AAATCTAGAATGACGGTGCTCGAAGAGATGCG-3 | Used to generate a mutated variant of M6 AopW1, deleting the first 175 amino acids |
| M6AopW1_EcoRI_R | 5'-AAAGAATTCTGGTTGATCCCCCGTCCG-3 | |
| M6AopW1_XbaI_F | 5'-AAATCTAGAATGCCTCTACAGTCCATT-3 | Used to generate a mutated variant of M6 AopW1, deleting the last 10 amino acids |
| dCt10M6AopW1_Eco_R | 5'-AAAGAATTCGGGATCCCAGAGGTCGATGCGA-3 | |
| M6AopW1_XbaI_F | 5'-AAATCTAGAATGCCTCTACAGTCCATT-3 | Used to generate a mutated variant of M6 AopW1, deleting the last 25 amino acids |
| M6dCt25AopW1_Eco_R | 5'-AAAGAATTCGCTTGCTGCTGCCGCTTT-3 | |

| | | |
|---|---|---|
| M6AopW1_XbaI_F | 5'-AAATCTAGAATGCCTCTACAGTCCATT-3 | Used to generate a mutated variant of M6 AopW1, deleting the last 50 amino acids |
| dCt50M6AopW1_Eco_R | 5'-AAAGAATTCGGACGCCGCACCCGAATT-3 | |
| M6AopW1_attB1 | 5'-GGGGACAAGTTTGTACAAAAAAGCAGGCTTAATGCCTCTACAGTTCATT-3 | Used for cloning of the M6 *aopW1* ORF without the stop codon in plasmid pDONR207 by the Gateway system |
| AopW1ns_attB2 | 5'-GGGGACCACTTTGTACAAGAAAGCTGGGTATGGTTGATCCCCCGTCCG-3 | |
| 7a1AopW1_attB1 | 5'-GGGGACAAGTTTGTACAAAAAAGCAGGCTTAATGCCTCTACAGTCCATT-3 | Used for cloning of the 7a1 *aopW1* ORF without the stop codon in plasmid pDONR207 by the Gateway system |
| AopW1ns_attB2 | 5'-GGGGACCACTTTGTACAAGAAAGCTGGGTATGGTTGATCCCCCGTCCG-3 | |
| 7a1AopW1cTP_attB1 | 5'-GGGGACAAGTTTGTACAAAAAAGCAGGCTTAATGGACCGGTGCATCCACAAG -3 | Used for cloning of the 7a1 *aopW1* ORF without first 255 bp (to eliminate the AopW1 chloroplast transit peptide or cTP) in plasmid pDONR207 by the Gateway system |
| AopW1ns_attB2 | 5'-GGGGACCACTTTGTACAAGAAAGCTGGGTATGGTTGATCCCCCGTCCG-3 | |
| 100NtAopW1pET-Eco_F | 5'-ATAGAATTCGCGCGCGACCAGGCAGCGGATTT-3 | Used to amplify the ORF of M6 and 7a1 *aopW1* without the first 300 bp for generation of recombinant AopW1 for *in vitro* actin depolymerization assays |
| AopW1_XhoI_R | 5'-ATACTCGAGTGGTTGATCCCCCGTCCGAGCA-3 | |

[1]For amplification by restriction-free cloning.

[2]Underlined nucleotides indicate the restriction sites of the corresponding enzymes in the primer names or the attB1 or attB2 sequences indicated in the primer names. Bolded nucleotides in primers used for mutagenesis of *aopW1* indicate mutated positions in forward and reverse primers.

**Supplementary Appendix S1.** *In silico* analysis of the AopW1 chloroplast localization.

AopW1 sequences used for *in silico* analysis: M6 (representative of group I) and AAC00-1 (representative of group II; identical to 7a1).

**ChloroP1.1 software** (Emanuelsson *et al*., 1999)

| Name | Length | Score | cTP* | CS-score | cTP-length |
|---|---|---|---|---|---|
| AopW1 (Ac M6) | 485 | 0.566 | Y | 4.430 | 59 |
| AopW1 (Ac AAC00-1) | 485 | 0.568 | Y | 4.430 | 59 |

* chloroplast transit peptides (cTP)
Y" means that the sequence *is* predicted to contain a cTP
"-" means that is predicted *not* to contain a cTP.

**WoLF PSORT software** (Horton *et al*., 2007)

***A. citrulli* M6 details. chlo: 12, cyto: 1, mito: 1**

PSORT features and traditional PSORTII prediction

14 Nearest Neighbors

| id | site | distance | identity | comments |
|---|---|---|---|---|
| STAD_RICCO | chlo | 234.839 | 15.2577% | [Uniprot] SWISS-PROT45:Chloroplast of green tissue and plastids of nonphotosynthetic tissues. |
| RAA3_CHLRE | chlo | 246.799 | 8.7493% | [Uniprot] SWISS-PROT45:Chloroplast stroma. |
| At3g22890.1 | chlo | 259.467 | 12.5% | [Arath] |
| STAD_CARTI | chlo | 260.851 | 15.4639% | [Uniprot] SWISS-PROT45:Chloroplast of green tissue and plastids of nonphotosynthetic tissues. |
| AROF_TOBAC | chlo | 267.692 | 13.2597% | [Uniprot] SWISS-PROT45:Chloroplast. |
| AROF_SOLTU | chlo | 271.298 | 12.616% | [Uniprot] SWISS-PROT45:Chloroplast. |

| | | | | |
|---|---|---|---|---|
| AROG_LYCES | chlo | 273.183 | 12.3616% | [Uniprot] SWISS-PROT45:Chloroplast. |
| RK29_MAIZE | chlo | 276.101 | 11.7526% | [Uniprot] SWISS-PROT45:Chloroplast. |
| NAC2_CHLRE | chlo | 281.515 | 11.4079% | [Uniprot] SWISS-PROT45:Chloroplast stroma. |
| GLYM_FLAPR | mito | 282.096 | 12.9094% | [Uniprot] SWISS-PROT45:Mitochondrial. |
| SR52_HORVU | cyto | 283.939 | 14.4531% | [Uniprot] SWISS-PROT45:Cytoplasmic. |
| HEM2_SPIOL | chlo | 285.462 | 13.963% | [Uniprot] SWISS-PROT45:Chloroplast. |
| STAD_CUCSA | chlo | 291.196 | 14.6392% | [Uniprot] SWISS-PROT45:Chloroplast of green tissue and plastids of nonphotosynthetic tissues. |
| AROF_ARATH | chlo | 292.069 | 12.3574% | [Arath] [Uniprot] SWISS-PROT45:Chloroplast. Evidence:TAS Pubmed:0012177500,1681544 |

***A. citrulli*** **AAC00-1** <u>**details**</u>**. chlo: 13, mito: 1**

PSORT features and traditional PSORTII prediction

14 Nearest Neighbors

| id | site | distance | identity | comments |
|---|---|---|---|---|
| RAA3_CHLRE | chlo | 217.735 | 9.1419% | [Uniprot] SWISS-PROT45:Chloroplast stroma. |
| STAD_RICCO | chlo | 228.148 | 15.0515% | [Uniprot] SWISS-PROT45:Chloroplast of green tissue and plastids of nonphotosynthetic tissues. |
| AROF_TOBAC | chlo | 249.771 | 12.9151% | [Uniprot] SWISS-PROT45:Chloroplast. |
| NAC2_CHLRE | chlo | 254.24 | 11.5523% | [Uniprot] SWISS-PROT45:Chloroplast stroma. |
| AROG_LYCES | chlo | 257.304 | 11.8299% | [Uniprot] SWISS-PROT45:Chloroplast. |
| AROF_SOLTU | chlo | 257.788 | 12.2677% | [Uniprot] SWISS-PROT45:Chloroplast. |

| | | | | |
|---|---|---|---|---|
| STAD_CARTI | chlo | 260.9 | 15.0515% | [Uniprot] SWISS-PROT45:Chloroplast of green tissue and plastids of nonphotosynthetic tissues. |
| HEM2_SPIOL | chlo | 267.28 | 13.347% | [Uniprot] SWISS-PROT45:Chloroplast. |
| MBB1_CHLRE | chlo | 274.186 | 16.0121% | [Uniprot] SWISS-PROT45:Chloroplast stroma. |
| RK29_MAIZE | chlo | 276.526 | 11.7526% | [Uniprot] SWISS-PROT45:Chloroplast. |
| ADT1_ARATH | mito | 276.997 | 16.701% | [Uniprot] SWISS-PROT45:Integral membrane protein. Mitochondrial inner membrane. |
| AROF_LYCES | chlo | 288.394 | 13.6452% | [Uniprot] SWISS-PROT45:Chloroplast. |
| STAD_CUCSA | chlo | 288.78 | 14.433% | [Uniprot] SWISS-PROT45:Chloroplast of green tissue and plastids of nonphotosynthetic tissues. |
| At4g27070.1 | chlo | 289.054 | 13.7014% | [Arath] |

**Localizer software** (Sperschneider *et al.*, 2017)

**A) Prediction for mature effector sequence without signal peptide.**

# -----------------
### LOCALIZER 1.0 Predictions
# -----------------

| Identifier | Chloroplast | Mitochondria | Nucleus |
|---|---|---|---|
| Ac M6 | Y (0.995 \| 6-33) | - | - |
| Ac AAC00-1 | Y (0.995 \| 6-33) | - | - |

### Proteins analyzed: 2
### Number of proteins with cTP: 2 (100.0%)
### Number of proteins with cTP & possible mTP: 0 (0.0%)
### Number of proteins with cTP & NLS: 0 (0.0%)
### Number of proteins with cTP & possible mTP & NLS: 0 (0.0%)
### Number of proteins with mTP: 0 (0.0%)
### Number of proteins with mTP & possible cTP: 0 (0.0%)
### Number of proteins with mTP & NLS: 0 (0.0%)
### Number of proteins with mTP & possible cTP & NLS: 0 (0.0%)

### Number of proteins with NLS and no transit peptides: 0 (0.0%)
### Summary statistics
### Number of proteins with chloroplast localization (cTP, cTP & possible mTP, cTP & NLS, cTP & possible mTP & NLS): 2 (100.0%)
### Number of proteins with mitochondrial localization (mTP, mTP & possible cTP, mTP & NLS, mTP & possible cTP & NLS): 0 (0.0%)
### Number of proteins with nuclear localization and no transit peptides: 0 (0.0%)
### Number of proteins with nuclear localization and with transit peptides: 0 (0.0%)

**B) Prediction for full sequence.**

# -----------------
### LOCALIZER 1.0 Predictions
# -----------------

| Identifier | Chloroplast | Mitochondria | Nucleus |
|---|---|---|---|
| Ac M6 | Y (0.985 \| 1-50) | - | - |
| Ac AAC00-1 | Y (0.978 \| 1-33) | - | - |

### Proteins analyzed: 2
### Number of proteins with cTP: 2 (100.0%)
### Number of proteins with cTP & possible mTP: 0 (0.0%)
### Number of proteins with cTP & NLS: 0 (0.0%)
### Number of proteins with cTP & possible mTP & NLS: 0 (0.0%)
### Number of proteins with mTP: 0 (0.0%)
### Number of proteins with mTP & possible cTP: 0 (0.0%)
### Number of proteins with mTP & NLS: 0 (0.0%)
### Number of proteins with mTP & possible cTP & NLS: 0 (0.0%)
### Number of proteins with NLS and no transit peptides: 0 (0.0%)
### Summary statistics
### Number of proteins with chloroplast localization (cTP, cTP & possible mTP, cTP & NLS, cTP & possible mTP & NLS): 2 (100.0%)
### Number of proteins with mitochondrial localization (mTP, mTP & possible cTP, mTP & NLS, mTP & possible cTP & NLS): 0 (0.0%)
### Number of proteins with nuclear localization and no transit peptides: 0 (0.0%)
### Number of proteins with nuclear localization and with transit peptides: 0 (0.0%)

**C) Prediction for full effector sequences with signal peptides.**

**C.1.) First 29 amino acids deleted.**

# -----------------
### LOCALIZER 1.0 Predictions
# -----------------

| Identifier | Chloroplast | Mitochondria | Nucleus |
|---|---|---|---|
| Ac M6 | Y (0.983 \| 38-62) | - | - |
| Ac AAC00-1 | Y (0.983 \| 38-62) | - | - |

### Proteins analyzed: 2
### Number of proteins with cTP: 2 (100.0%)
### Number of proteins with cTP & possible mTP: 0 (0.0%)
### Number of proteins with cTP & NLS: 0 (0.0%)
### Number of proteins with cTP & possible mTP & NLS: 0 (0.0%)
### Number of proteins with mTP: 0 (0.0%)
### Number of proteins with mTP & possible cTP: 0 (0.0%)
### Number of proteins with mTP & NLS: 0 (0.0%)
### Number of proteins with mTP & possible cTP & NLS: 0 (0.0%)
### Number of proteins with NLS and no transit peptides: 0 (0.0%)
### Summary statistics
### Number of proteins with chloroplast localization (cTP, cTP & possible mTP, cTP & NLS, cTP & possible mTP & NLS): 2 (100.0%)
### Number of proteins with mitochondrial localization (mTP, mTP & possible cTP, mTP & NLS, mTP & possible cTP & NLS): 0 (0.0%)
### Number of proteins with nuclear localization and no transit peptides: 0 (0.0%)
### Number of proteins with nuclear localization and with transit peptides: 0 (0.0%)

**C.2.) First 30 amino acids deleted.**

# -----------------
### LOCALIZER 1.0 Predictions
# -----------------

| Identifier | Chloroplast | Mitochondria | Nucleus |
|---|---|---|---|
| Ac M6 | - | - | - |
| Ac AAC00-1 | - | - | - |

### Proteins analyzed: 2
### Number of proteins with cTP: 0 (0.0%)
### Number of proteins with cTP & possible mTP: 0 (0.0%)
### Number of proteins with cTP & NLS: 0 (0.0%)
### Number of proteins with cTP & possible mTP & NLS: 0 (0.0%)
### Number of proteins with mTP: 0 (0.0%)
### Number of proteins with mTP & possible cTP: 0 (0.0%)
### Number of proteins with mTP & NLS: 0 (0.0%)
### Number of proteins with mTP & possible cTP & NLS: 0 (0.0%)
### Number of proteins with NLS and no transit peptides: 0 (0.0%)
### Summary statistics
### Number of proteins with chloroplast localization (cTP, cTP & possible mTP, cTP & NLS, cTP & possible mTP & NLS): 0 (0.0%)
### Number of proteins with mitochondrial localization (mTP, mTP & possible cTP, mTP & NLS, mTP & possible cTP & NLS): 0 (0.0%)
### Number of proteins with nuclear localization and no transit peptides: 0 (0.0%)
### Number of proteins with nuclear localization and with transit peptides: 0 (0.0%)
